# Separate physiological roles of specific and non-specific DNA binding of HU protein in *Escherichia coli*

**DOI:** 10.1101/2021.06.17.448862

**Authors:** Subhash Verma, Sankar Adhya

**Affiliations:** Laboratory of Molecular Biology, Center for Cancer Research, National Cancer Institute, Bethesda, Maryland 20892

## Abstract

Conserved in bacteria, the histone-like protein HU is crucial for genome organization and expression of many genes. It binds DNA regardless of the sequence and exhibits two binding affinities in vitro, low-affinity to any B-DNA (non-specific) and high-affinity to DNA with distortions like kinks and cruciforms (structure-specific), but the physiological relevance of the two binding modes needed further investigation. We validated and defined the three conserved lysine residues, K3, K18, and K83, in *Escherichia coli* HU as critical amino acid residues for both non-specific and structure-specific binding and the conserved proline residue P63 additionally for only the structure-specific binding. By mutating these residues in vivo, we showed that two DNA binding modes of HU play separate physiological roles. The DNA structure-specific binding, occurring at specific sites in the *E. coli* genome, promotes higher-order DNA structure formation, regulating the expression of many genes, including those involved in chromosome maintenance and segregation. The non-specific binding participates in numerous associations of HU with the chromosomal DNA, dictating chromosome structure and organization. Our findings underscore the importance of DNA structure in transcription regulation and promiscuous DNA-protein interactions in a dynamic organization of a bacterial genome.

## INTRODUCTION

The *Escherichia coli* genome of 1.5-millimeter contour length is condensed and organized into a helical ellipsoid of ∼1.0 µ^3^ volume that together with its associated RNAs and proteins constitute the *E. coli* chromosome (nucleoid) (Fisher et al., 2013; Verma et al., 2019). The concerted action of the intrinsic DNA properties, and many biochemical factors leads to the formation of the chromosome (Verma et al., 2019). Among the biochemical factors, the histone-like protein HU and the structural maintenance of chromosomes (SMC) complex MukBEF have recently emerged as crucial players for the genome organization as a whole (Lioy et al., 2018; Makela and Sherratt, 2020b; Walker et al., 2020). HU also facilitates gene expression (Prieto et al., 2012) and gene regulatory processes (Aki and Adhya, 1997), chromosomal and prophage DNA replication (Hwang and Kornberg, 1992; Montano et al., 2012; Ogura et al., 1990), DNA repair (Kamashev and Rouviere-Yaniv, 2000), as well as site-specific recombination (Zablewska and Kur, 1995). Additionally, HU regulates bacterial motility (Nishida et al., 1997). Thus, HU plays a major role in bacterial physiology. Not surprisingly, HU is ubiquitous in bacterial kingdom and essential in many bacterial species including many pathogens (Bhowmick et al., 2014; Fernandez et al., 1997). Composed of two homologous subunits α and β encoded by *hupA* and *hupB* genes respectively, the *E. coli* HU can exist as three dimeric forms: Huα_2_, Huβ_2_, and HUαβ. The composition of HU varies according to growth cycle, with HUα_2_ being predominant in the exponential phase and HUαβ being predominant in the stationary phase while HUβ_2_ is nearly undetectable in any phase (Claret and Rouviere-Yaniv, 1997).

HUα_2_ and HUαβ bind DNA independent of base pair sequence but exhibits two distinct DNA binding affinities in vitro. They bind with low-affinity to B-form dsDNA but bind with a 100-fold higher affinity to distorted dsDNA with kinks, nicks, gaps, or cruciform structure (Bonnefoy et al., 1994; Kamashev and Rouviere-Yaniv, 2000; Pinson et al., 1999; Pontiggia et al., 1993; Vitoc and Mukerji, 2011). Hereafter, we refer the low-affinity binding as DNA *non-specific* binding and the high-affinity binding to distorted DNA as DNA *structure-specific* binding. HU is a Y-shaped dimer wherein two homo- or heteromeric subunits intertwine to form a α-helical “body” and two extending β-strands or “arms” (Guo and Adhya, 2007; Swinger et al., 2003). In crystal structures of *E. coli* Huα_2_ or HUαβ bound to a 20-bp DNA of random nucleotide sequence (Hammel et al., 2016), mostly three surface exposed lysine residues (K3, K18, and K83) effect non-specific binding through electrostatic interactions with the DNA phosphate backbone (phosphate locks). The lysine residues are present on either side of the dimer allowing a single dimer to interact with two DNA segments through each subunit. The non-specific binding triggers multimerization of HUα_2_ dimers, but not of HU dimers, resulting in three-dimensional intermolecular DNA *bundling* (Hammel et al., 2016). Controlled by ionic strength and pH, the intermolecular hydrogen bonds between residues E34K and K37 dictates the HUα_2_ multimerization, potentially functioning as a molecular switch that regulates DNA bundling in an environment dependent manner (Hammel et al., 2016; Remesh et al., 2020).

In crystal structures of HU in a complex with a distorted dsDNA with kinks, a highly conserved proline residue at position 63 (P63) is critical for the structure-specific binding (Guo and Adhya, 2007; Swinger et al., 2003). While the positively charged surface of the two β rms reach around the opposite faces of the DNA wrapping the minor grooves, the P63 residues at the tip of each β-arm intercalates into bases in the corresponding minor groove. The two kinks, which are 9 bp apart, coincide with the intercalation of P63 residues, suggesting that P63 intercalation induce and/or stabilize the kinks. HU binds using the same structural motif of paired inclined helices to other forms of distorted DNA such as cruciform and nicked DNA (Balandina et al., 2002; Kamashev et al., 1999).

The contributions, if any, of the lysine residues in DNA structure-specific binding, or that of the P63 residue in non-specific binding are so far unresolved. The DNA in the structure-specific complex in the crystal structures was not long enough to allow contacts with the lysine residues on the sides of the HU “body” (Swinger et al., 2003). Also, the electron density of the β-arms was not visible in the crystal structures of HU with the 20 bp dsDNA, implying that the P63 residue was not important for the non-specific binding (Hammel et al., 2016), except that P63 residue may stabilize the intermolecular DNA bundling through β-arm-β-arm interactions via the formation of a β-zipper (Remesh et al., 2020). Despite the extensive in vitro characterization, the details of the two DNA binding modes in the HU-dependent chromosome structure and function in vivo remains to be investigated. In this study, we determined specific contribution of different amino acid residues in the structure-specific and the non-specific DNA binding of HUα_2_ in variously proposed physiological processes both in vitro and in vivo. We found that processes needing structure-specific binding require both P63 and lysine residues of HUα_2_, but those involving non-specific DNA binding use only the lysine residues corroborating the structural studies. The requirement of both the lysine residues and P63 residue in specific binding provides a more detailed mechanism of structural-specific binding to a distorted DNA in which the β-arm and the P63 residue bend the DNA further because of the lysine mediated phosphate locks further along the DNA on each side. The structure-specific binding, while not necessary for overall chromosome structure, is critical for the regulation of specific gene expressions by HU. On the contrary, HU binds to DNA as part of the chromosome structure primarily through non-specific binding as we previously suggested from single cell analysis (Bettridge et al., 2020). Our findings reveal distinct physiological roles of both structure-specific and non-specific DNA binding modes of HU.

## RESULTS

### Amino acid residues of HU involved in specific and non-specific DNA binding in vitro

The P63 residue in the β-arm of HU is involved in structure-specific binding (Guo and Adhya, 2007; Swinger et al., 2003). To test any additional role of the three conserved lysine residues (K3, K18, and K83) required for non-specific binding in the structure-specific DNA binding, we used purified wild-type HUα_2_ and HUα_2_ with P63A mutation, K3A, K83 double mutations, or K3A, K18A, KA83 triple mutations to follow HU binding to an artificially designed cruciform DNA (Vitoc and Mukerji, 2011) and a 40-bp double-stranded DNA of random sequence. Binding was measured by fluorescence polarization in solution; the reactions were further loaded onto electrophoretic gels to measure the formation of discrete DNA-protein complexes by electrophoretic mobility shift assays (EMSA).

We observed cooperative binding of wild-type HUα_2_ to both the linear duplex and cruciform DNA in solution with similar calculated Hill coefficients (*h*) and dissociation constants (*K_d_)* (non-specific binding: *h* = 3.2 and *K_d_* = 0.101 µM; structure-specific binding: *h* = 2.8 and *K_d_* = 0.096 µM) (Figure 1). However, the linear duplex did not migrate as discrete retarded species in EMSA, consistent with weak and transitory non-specific binding (Bettridge et al., 2020; Pinson et al., 1999), but the cruciform DNA migrated as four discrete bands, indicative of strong, cooperative binding of each cruciform DNA molecule to four molecules of HUα_2_ as suggested previously (Bonnefoy et al., 1994). The P63A mutation reduced the cooperative binding to cruciform DNA with *K_d_ =*0.149 µM, *h*=1.6, validating the importance of P63 in binding to a distorted DNA as suggested by crystal structures of HU-structure-specific DNA complexes (Guo and Adhya, 2007; Swinger et al., 2003), but it did not affect the binding to the linear DNA duplex (*K_d_* 0.106 µM, *h* = 3.0). In contrast, the K3A, K83A double and K3A, K18A, K83A triple mutations not only reduced the cooperative binding to linear duplex, as expected, they also significantly reduced the same to cruciform DNA (Figures 1). These results suggest that beside P63, the lysine residues are also critical for the structure-specific binding of HU, but P63 does not participate in non-specific binding.

**Figure 1.**
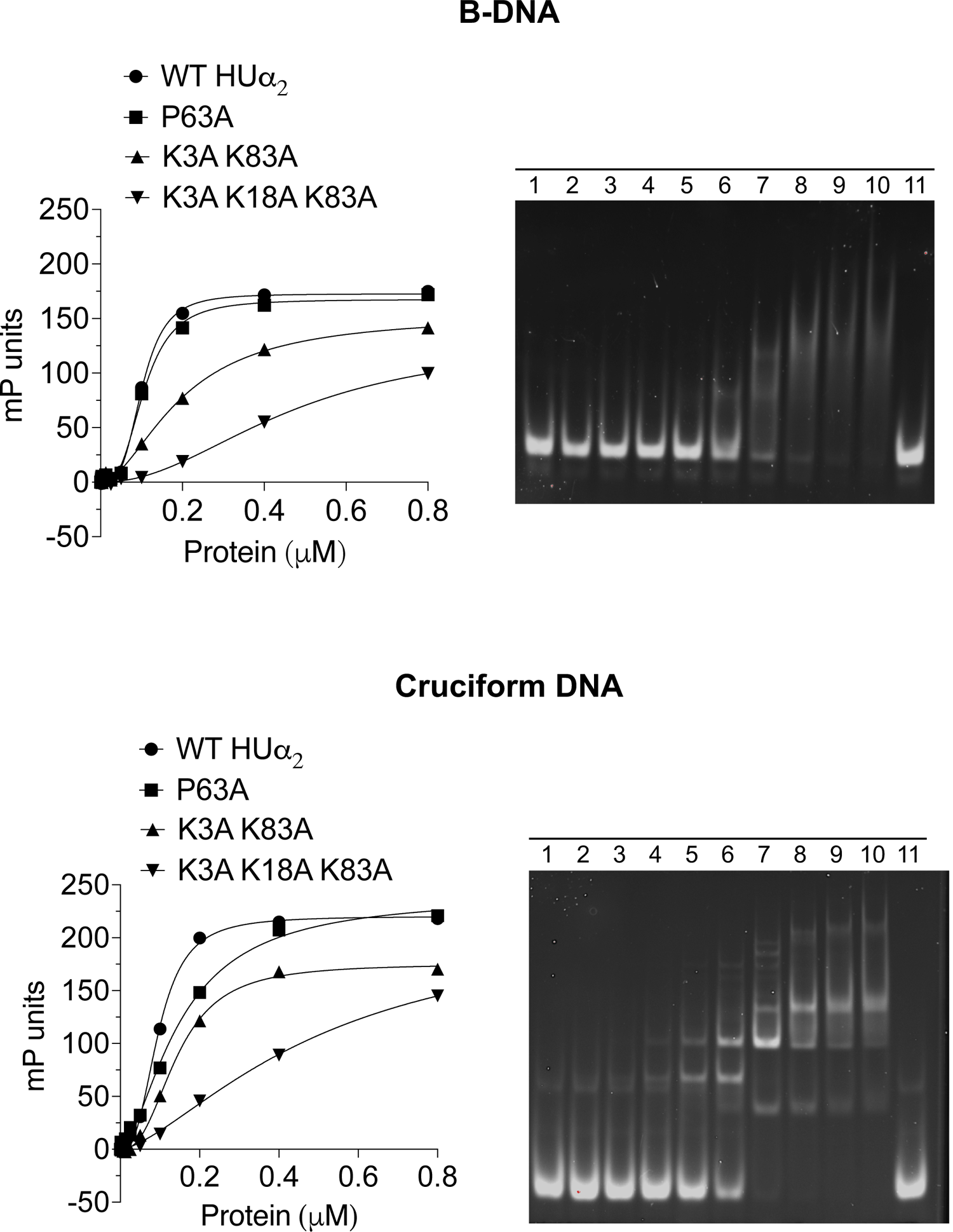
Amino acid residues of HU involved in specific and non-specific DNA binding in vitro. Binding of purified wild-type HUα_2_ or its variants to a 6-carboxy fluorescein labeled 40-bp double stranded DNA of random sequence (B-DNA) or 40-bp artificially designed cruciform DNA. On the left, each data point on the Y-axis represents the milipolarization (mP) units measured by fluorescence polarization. The line represents the best fit of the data points using the hill slope equation. On the right, the binding reactions of wild-type HUα_2_ analyzed by electrophoretic mobility shift assay. Lanes 1 to 10 contain 0.01, 0.03, 0.06, 0.12, 0.25, 0.5, 0.1, 0.2, 0.4, and 0.8 µM wild-type HUα_2_. Lane 11 contains only the DNA.

### Involvement of structure-specific binding of HU in DNA supercoil restraining

HU constrains DNA when supercoiled (Guo and Adhya, 2007). The crystal structures of HU with a distorted (artificially bent) DNA suggest that HU forms negative supercoils by inducing/stabilizing two kinks in DNA. The kinks are 9 bp apart with bend angles of opposite angular orientations, mimicking a negative writhe (Guo and Adhya, 2007; Swinger et al., 2003), suggesting that the negative writhe may be responsible for HU-mediated supercoil retention in DNA. The intercalation of stacked bases by the P63 residues in the two subunits coincides with the site of kinks, indicating that P63 induces and/or stabilizes the kinks. We tested the importance of P63 and the lysine residues for supercoil restraining ability of HU by incubating the wild-type HUα_2_ and HUα_2_ variant proteins with a relaxed plasmid DNA, made by introducing a nick in one of the strands. After incubation, the nicks were sealed with DNA ligase, the proteins removed, and the resulting DNA topoisomers were resolved on gels by EMSA. We observed the appearance of supercoils at 0.4 µM with saturation at 1.6 µM concentrations with wild type HUα_2_ (Figure 2). The P63A mutation significantly reduced the appearance of HU dependent supercoils even at the saturating levels of the mutant protein, demonstrating a role of structure-specific binding of HUα_2_ in DNA supercoils. We also found the lysine residues played some roles in the process because the K3A-K83A double mutations in HUα_2_ also reduced the ability of helping supercoil formation (Figure 2). From the results of DNA binding described above together with the supercoiling assays, we conclude that the DNA structure-specific binding of HU induces negative supercoils in DNA.

**Figure 2.**
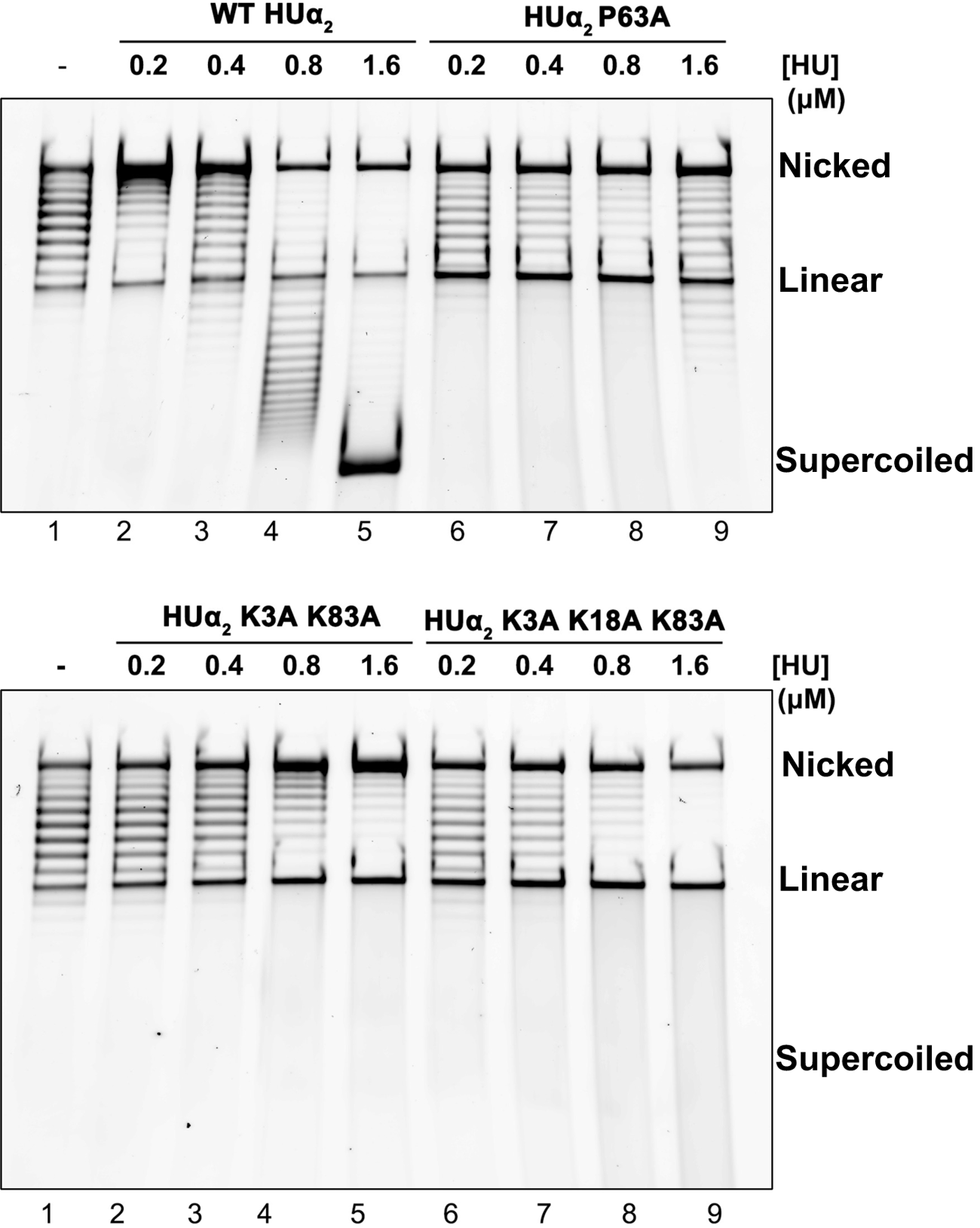
Involvement of structure-specific binding of HU in DNA supercoil restraining. The ability of wild-type HUα_2_ protein or its variants to induce negative supercoils in singly nicked plasmid DNA. A 11090-bp singly nicked plasmid was incubated with the proteins at indicated concentrations and then resealed using T4 DNA ligase. After removing the protein, the DNA was resolved on the 0.8% agarose gel for 18 hours and stained with SYBER gold.

### Structure-specific binding in specific gene expression: the gal operon

HU binds specifically to the apical region of the DNA segment between two operator sites of the GalR repressor at the *gal* operon in the *E. coli* chromosome (Majumdar and Adhya, 1984). The binding promotes the formation of a DNA loop by DNA-bound GalR-GalR interactions that represses the transcription from the *P2* promoter located within the looped DNA (Lyubchenko et al., 1997). HU binding stabilizes the DNA bending at the apex, occurring due to transient GalR-GalR interactions (Geanacopoulos et al., 2001). The HU binding requires the formation of a cruciform DNA structure generated at the apex (HU binding site or *hbs*) by DNA supercoiling (Figure S1) (Lewis et al., 1999; Lia et al., 2003). The *gal* system has served as a paradigm for the study of DNA structure-specific binding by HU in gene regulatory systems. We tested the importance of P63 and the lysine residues in the *gal* system by examining the ability of purified wild-type and variant HUα_2_ proteins to repress *in vitro* transcription from the DNA looping-sensitive *P2* promoter in a supercoiled DNA template (Lewis et al.,1999). We observed that, compared to the wild-type, the P63A mutation alone dramatically reduced the ability of HUα_2_ to repress the transcription (Figures 3A and S2), confirming a critical role of P63 in the cruciform DNA binding in a physiologically relevant system. Similarly, the K3A-K83A and K3A-K18A-K83A mutations also reduced the ability of HUα_2_ to repress the transcription (Figures 3A and S2), confirming that lysine residues also contribute to the binding of the cruciform structure.

**Figure 3.**
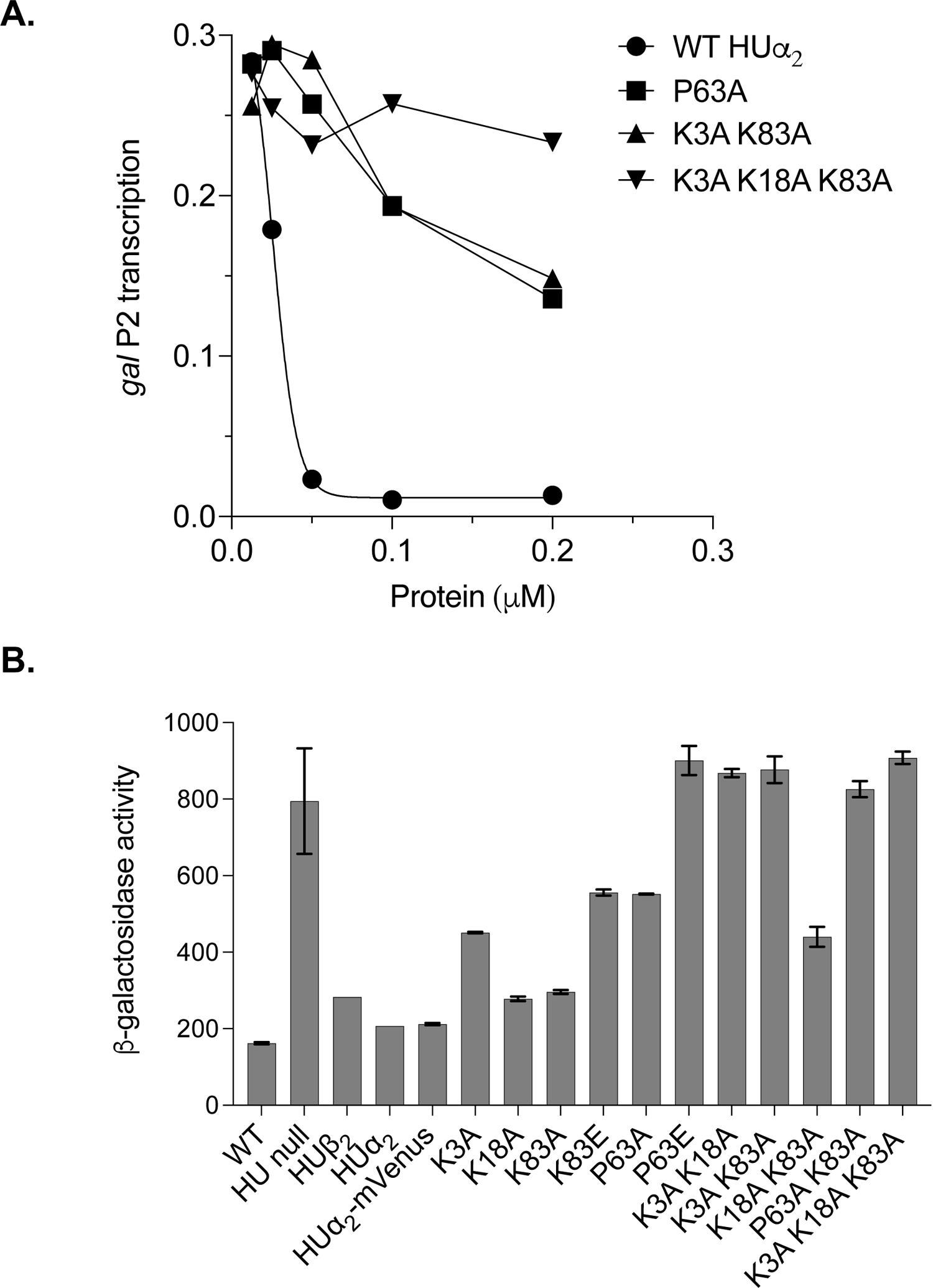
The role of structure-specific DNA binding of HU in the transcription of galactose utilization genes A. The in vitro transcription from the DNA lopping sensitive *galP2* promoter in the presence of 200 nM GalR and wild-type HUα_2_ protein or its variants. Each data point represents the intensity of the *galP2* transcript band in the gel in Figure S2 after normalizing by the RNA 1 transcript band intensity in the same lane. **B.** β-galactosidase activity of the *gal-lacZ* transcription reporter in *E. coli* strains harboring wild-type *hup genes* (WT), Δ*hupA* Δ*hupB* alleles (HU null), Δ*hupA* allele producing only HUβ_2_, and Δ*hupB* allele producing only HUα_2_, HUα_2_-mVenus, or HUα_2_-mVenus with indicated mutations.

We also tested the importance of P63 and the lysine residues in the HU-mediated *gal* repression *in vivo.* We replaced the promoter of the *lac* operon in the chromosome with the *gal* promoter segment to create a *gal-lacZ* transcription reporter fusion. We introduced different mutations in the native *hupA* gene that encodes the α subunit and deleted the Δ*hupB* gene that encodes the β subunit. HUα_2_ was as capable of repressing the transcription from the *gal* promoter as the HUαβ heterodimer since the repressed levels of β-galactosidase activity were similar in the strains with wild-type *hup* genes and with Δ*hupB* allele producing only HUα_2_ (Figure 3B). Deleting both *hupA* and *hupB* genes resulted in an increase in the levels of β-galactosidase activity, consistent with the de-repression of the *P2* promoter due to the absence of the DNA loop (Lewis et al., 1999). The P63A mutation in the *hupA* gene caused nearly 3-fold increase in the levels of β-galactosidase activity, and a P63E substitution completely abolished the repression by HUα_2_ (Figure 3B), confirming the in vitro results that P63 is a critical amino acid residue in HUα_2_ for binding to DNA and to promote the formation of the DNA loop at the *gal* operon. When we mutated the lysine residues individually, the K3A and K83E substitutions showed de-repression similar to that of the P63A, and, in combinations, K3A-K18A, K3A-K83A, K18A-K83A, K3A-K18A-K83A mutations all showed complete or near complete de-repression (Figure 3B) supporting the in vitro finding about the contribution of the lysine residues in binding to the *gal* cruciform DNA in vivo.

### Structure-specific binding also regulate many other genes

After confirming the role of structure-specific binding of HU in the *gal* operon regulation by a DNA loop formation, we investigated the role of structure-specific binding in regulation of other genes in the *E. coli* chromosome. We analyzed transcriptome of the *E. coli* strains harboring wild-type *hup genes,* Δ*hupA* Δ*hupB* alleles, Δ*hupB allele* producing only HUα_2_, or *hupAP63A* Δ*hupB* alleles producing the HUα_2_ P63A mutant protein. After mapping the RNA reads to the genome, we followed the workflow described by Chen Y. et al (Chen et al., 2016) to identify any differentially expressed genes (DEGs). We used false discovery rate of ≤ 0.5% and log_2_ fold change ≥ 1.5 or ≤ −1.5 to define differential gene expression. Δ*hupB* strain producing HUα_2_ homodimer showed differential expression of only two genes (Figure 4) compared to the wild-type strain (WT) that most likely produces HUαβ heterodimers under the experimental growth conditions (Figure S3). These results suggested that HUα_2_ was as capable in controlling gene *hupB* strain producing no HU when compared to wild-type strain (Figure 4 and Table S2). 203 genes showed higher expression (up-regulation) and 235 genes showed lower expression (down-regulation).

**Figure 4.**
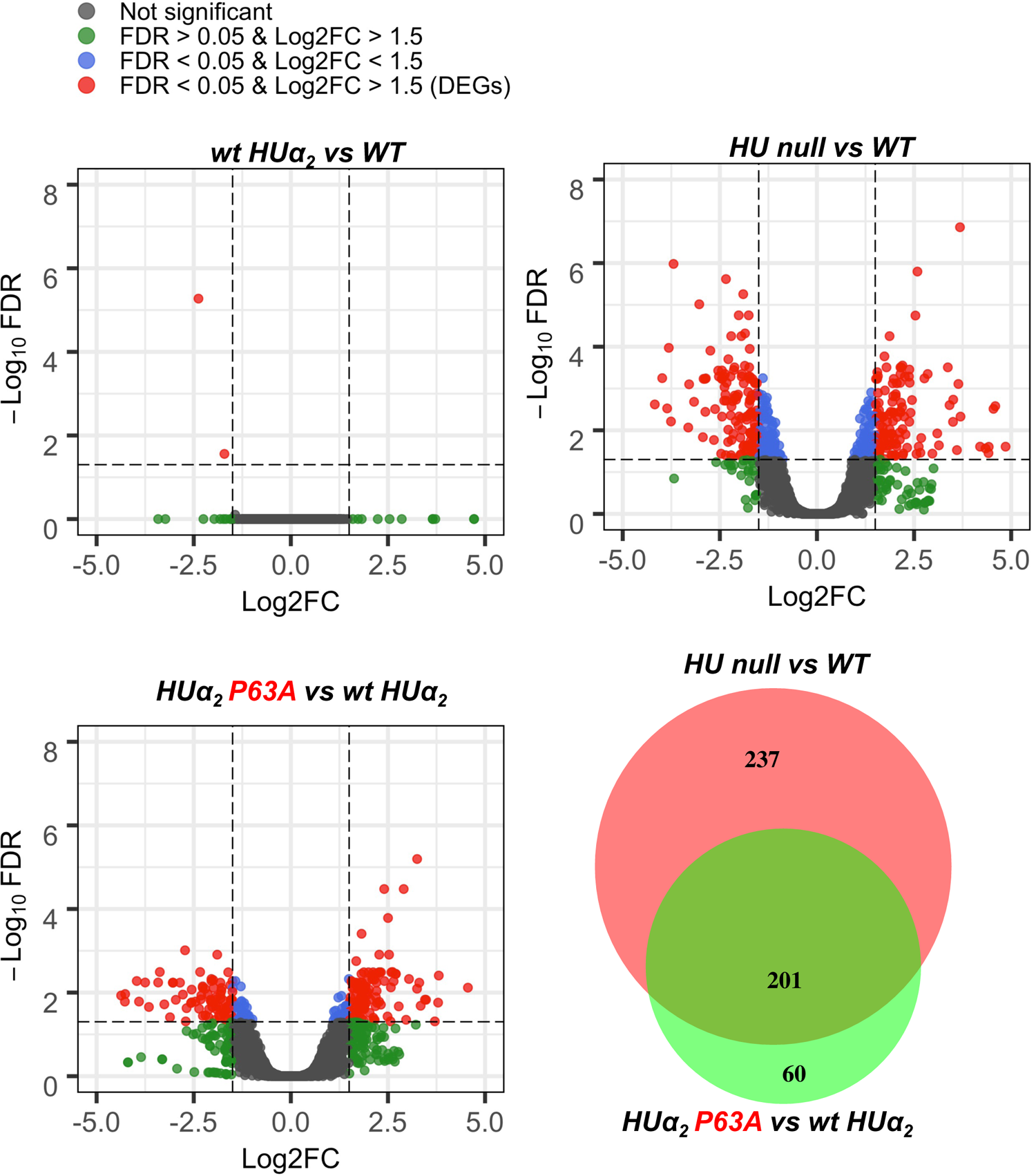
Involvement of structure-specific binding of HU in global gene expression. Volcano plots showing differential gene expression in the *E. coli* strain harboring Δ*hupA* Δ*hupB* alleles (HU null) compared to the strain harboring wild-type *hup* genes (WT), in the strain harboring Δ*hupB* allele producing only HUα_2_ (wt HUα_2_) compared to WT, and in the strain harboring ΔhupB *hupA P63A-mVenus* allele producing HUα_2_ P63A-mVenus (HUα_2_ P63A) Δ*hupB* allele producing only HUα_2_. y-axis represents inverse of Log10 value of false discovery rate (FDR), and x-axis represents Log2 value of the fold-change (Log2FC) in the transcript abundance. Horizontal dotted line represents 0.05 FDR cut off and vertical dotted line represents 1.5 log2FC cut off. DEGs denotes differentially expressed genes. Venn diagram showing overlap in DEGs between HU null versus WT and HUα_2_ P63A vs wt HUα_2_ conditions.

To measure any role of structure-specific binding of HU in DEGs, we compared gene expression between the strains expressing HUα_2_ and HUα_2_ P63A mutant protein. The strain expressing HUα_2_ P63A showed differential expression of 261 genes compared to the strain expressing HUα_2_ (Figure 4 and Table S3). 201 DEGs were common in both the HU-deleted strain and in the strain expressing HU P63A, and they showed differential expression in the same direction, up- or down-regulation, in both strains (Figure 4 and Table S4). These results show that the loss of the HU structure-specific binding is involved in the regulation of these genes. Since the P63A point mutation does not completely disrupt the structure-specific binding (Figure 1), we predicted the remaining genes may not have enough differential gene expression to be detected as DEGs by our criteria, but their differential expression would trend towards the HU-deleted strain. To test the prediction, we constructed the heatmap of normalized read counts (counts per million) for 438 DEGs identified in the HU-deleted strain (Figure 5). We found that the overall expression pattern of these genes in both the HU-deleted strain and in the strain expressing HUα_2_ P63A was similar, supporting the prediction that the complete disruption of the binding may account for even more DEGs observed in the HU-deleted host.

**Figure 5.**
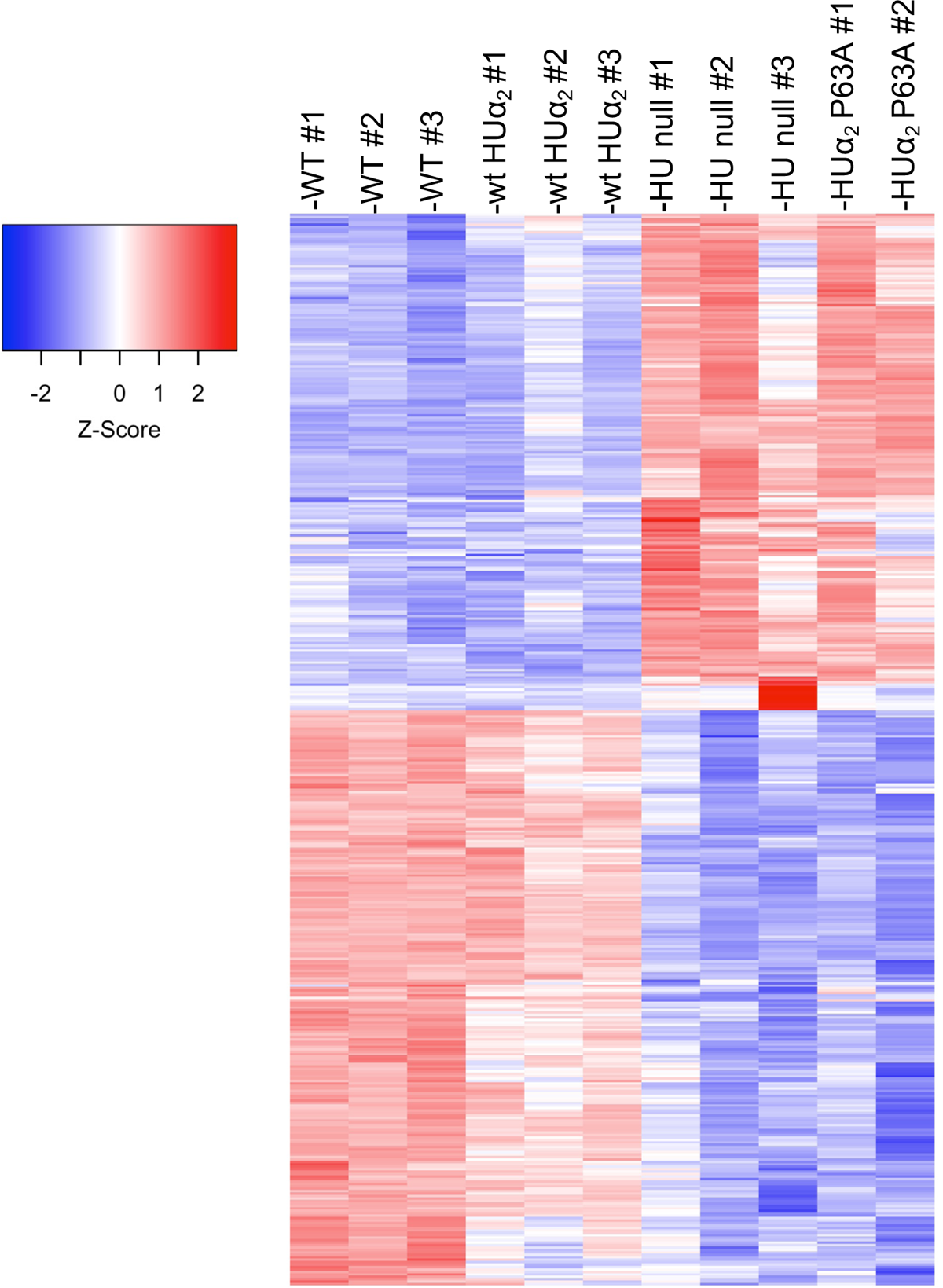
Overall expression pattern of genes identified as differentially expressed genes in the HU null strain. Heat map of normalized reads counts (counts per million) of 438 genes identified as differentially expressed genes in the HU null strain. Each column represents a single replicate of the strain indicated. Each row represents a single gene.

### Involvement of HU in bacteriophage replication

HU is important for the growth of bacteriophages Mu and P1 – Mu during its vegetative growth in replicative transposition, and P1 during replication in its prophage (plasmid) state (Montano et al., 2012; Ogura et al., 1990). It is known that HU binds to a specific region of Mu DNA at the left end of the prophage genome and promotes the assembly of a higher-order Mu transpososome structure by bending the DNA and/or stabilizing a pre-bent DNA (Montano et al., 2012). The Mu transpososome assembly triggers repeated replicative transposition of Mu DNA needed for phage growth in the *E. coli* host chromosome.

To understand what kind of HU binding occurs in Mu transpososome, we measured plaque-forming efficiency of Mu phage on *E. coli* lawns expressing either HUα_2_ or HUα_2_ with the mutations (Table 1). The efficiency of plating on the lawns of the *E. coli* strains expressing HUα_2_ with K3A or P63A mutation was about one order of magnitude lower than the strain expressing wild-type HUα_2_. But the combination of both mutations rendered the *E. coli* host completely resistant to the Mu growth. These results suggest that HU binding to Mu DNA involves a structure-specific binding involving both P63 and the lysine residues. We also measured the transformation efficiency of the *E. coli* strains expressing wild-type HUα_2_ or HUα_2_ with mutations using the P1 plasmid DNA, which is a measure of PI prophage replication. Compared to the wild-type strain expressing HUα_2_, we did not observe any transformants with *E. coli* expressing HUα_2_ with the K3A or the P63A mutation (Table 2) suggesting that the structure-specific binding of HU to P1 DNA is an absolute requirement for P1 plasmid replication.

**Table 1.** Effect of various mutations in HU on the Mu phage growth.

| Strain | Genotype | <sup>2</sup>PFU/ml |
| --- | --- | --- |
| MG1655 | WT | $6.6 \times 10^8$ |
| SCV129 | HU$\alpha_2$ | $6.3 \times 10^8$ |
| SCV130 | HU$\alpha_2$-*mVenus*<sup>1</sup> | $5.3 \times 10^8$ |
| SCV142 | K3A | $5.7 \times 10^7$ |
| SCV140 | K18A | $3.2 \times 10^8$ |
| SCV108 | K83A | $3.2 \times 10^8$ |
| SCV106 | P63A | $2.1 \times 10^7$ |
| SCV111 | K3A K83A | 0 |
| SCV133 | K83A P63A | 0 |
| SCV144 | K3A K18A K83A | 0 |

<sup>1</sup> Mutations indicated are in HU $\alpha_2$ -*mVenus*.
<sup>2</sup> Plaque forming units.

**Table 2.**
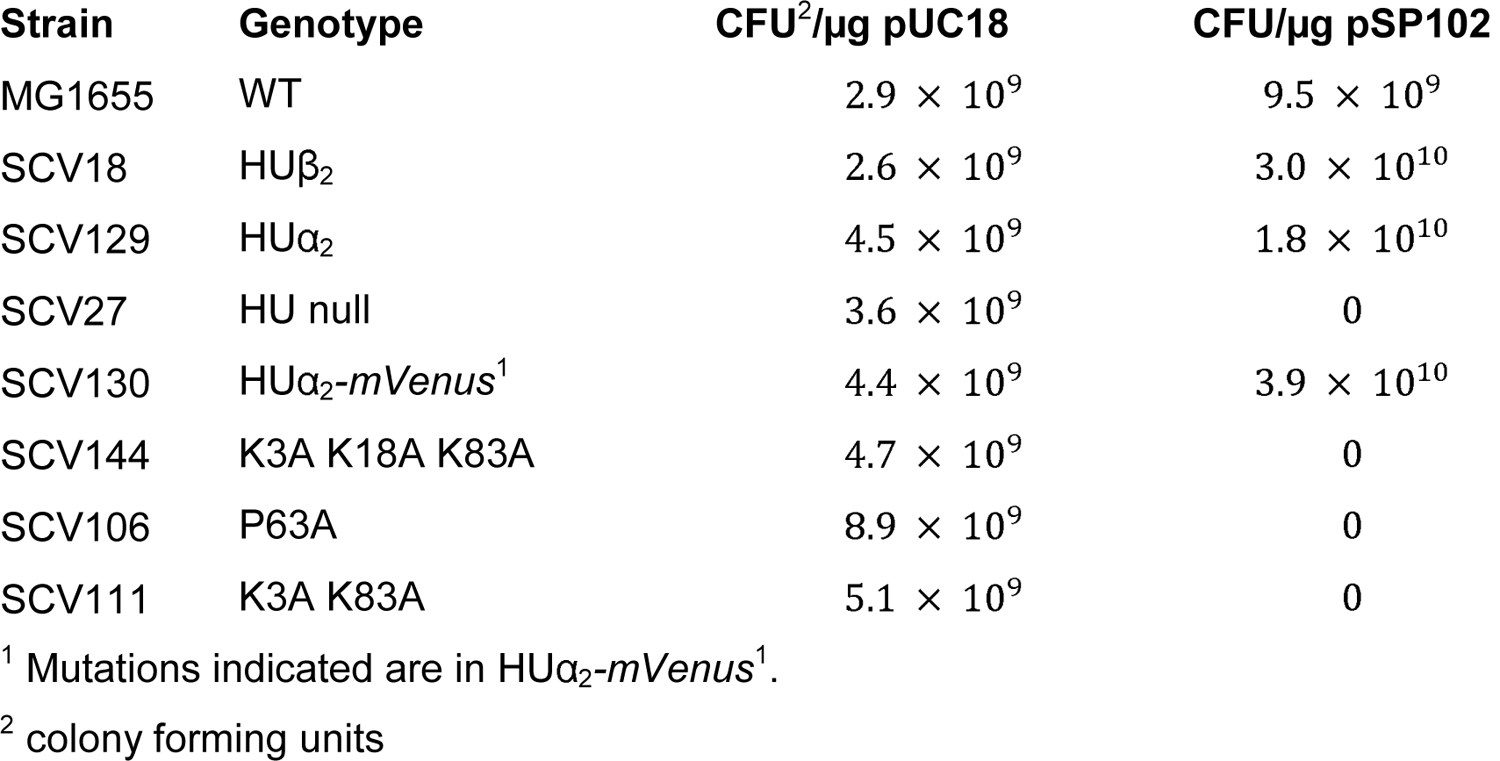
Effect of various mutations in HU on the P1 phage (plasmid state) growth.

| Strain | Genotype | CFU<sup>2</sup>/μg pUC18 | CFU/μg pSP102 |
| --- | --- | --- | --- |
| MG1655 | WT | $2.9 \times 10^9$ | $9.5 \times 10^9$ |
| SCV18 | HU$\beta_2$ | $2.6 \times 10^9$ | $3.0 \times 10^{10}$ |
| SCV129 | HU$\alpha_2$ | $4.5 \times 10^9$ | $1.8 \times 10^{10}$ |
| SCV27 | HU null | $3.6 \times 10^9$ | 0 |
| SCV130 | HU$\alpha_2$-*mVenus*<sup>1</sup> | $4.4 \times 10^9$ | $3.9 \times 10^{10}$ |
| SCV144 | K3A K18A K83A | $4.7 \times 10^9$ | 0 |
| SCV106 | P63A | $8.9 \times 10^9$ | 0 |
| SCV111 | K3A K83A | $5.1 \times 10^9$ | 0 |

<sup>1</sup> Mutations indicated are in HU $\alpha_2$ -*mVenus*<sup>1</sup>.
<sup>2</sup> colony forming units

### Nature of HU DNA binding in E. coli chromosome structure and function

Like HU involvement in the replicative processes of bacteriophage and plasmid chromosomes, HU has also been reported to influence replication and partitioning of *E. coli* chromosome (Bahloul et al., 2001; Chodavarapu et al., 2008; Dixon and Kornberg, 1984; Jaffe et al., 1997). We investigated whether HU contributes to such processes by DNA binding. First, we examined any impact of mutating P63 and the lysine residues of HUα_2_ on its chromosomal association, which is responsible of chromosomal structural maintenance (Lioy et al., 2018; Walker et al., 2020), by following the co-localization of HUα_2_ and its variants with the chromosome in vivo in single cells using wide-field fluorescence microscopy. Whereas most of the wild-type HUα_2_ and its P63A mutant co-localized very well with the chromosome with a distribution pattern resembling the shape of the chromosome, the HUα_2_ with the triple lysine mutations K3A-K18A-K83A did not co-localize with DNA in the chromosome (Figures 6A and 6B). These results, together with in vitro DNA binding assays described above (Figure 1), suggest that non-specific DNA binding, which requires only the conserved lysine residues but not P63, is the primary mode of overall HU-DNA interactions in the chromosome. These results do not rule out any role of structure-specific bindings at some limited sites in the chromosome.

**Figure 6.**
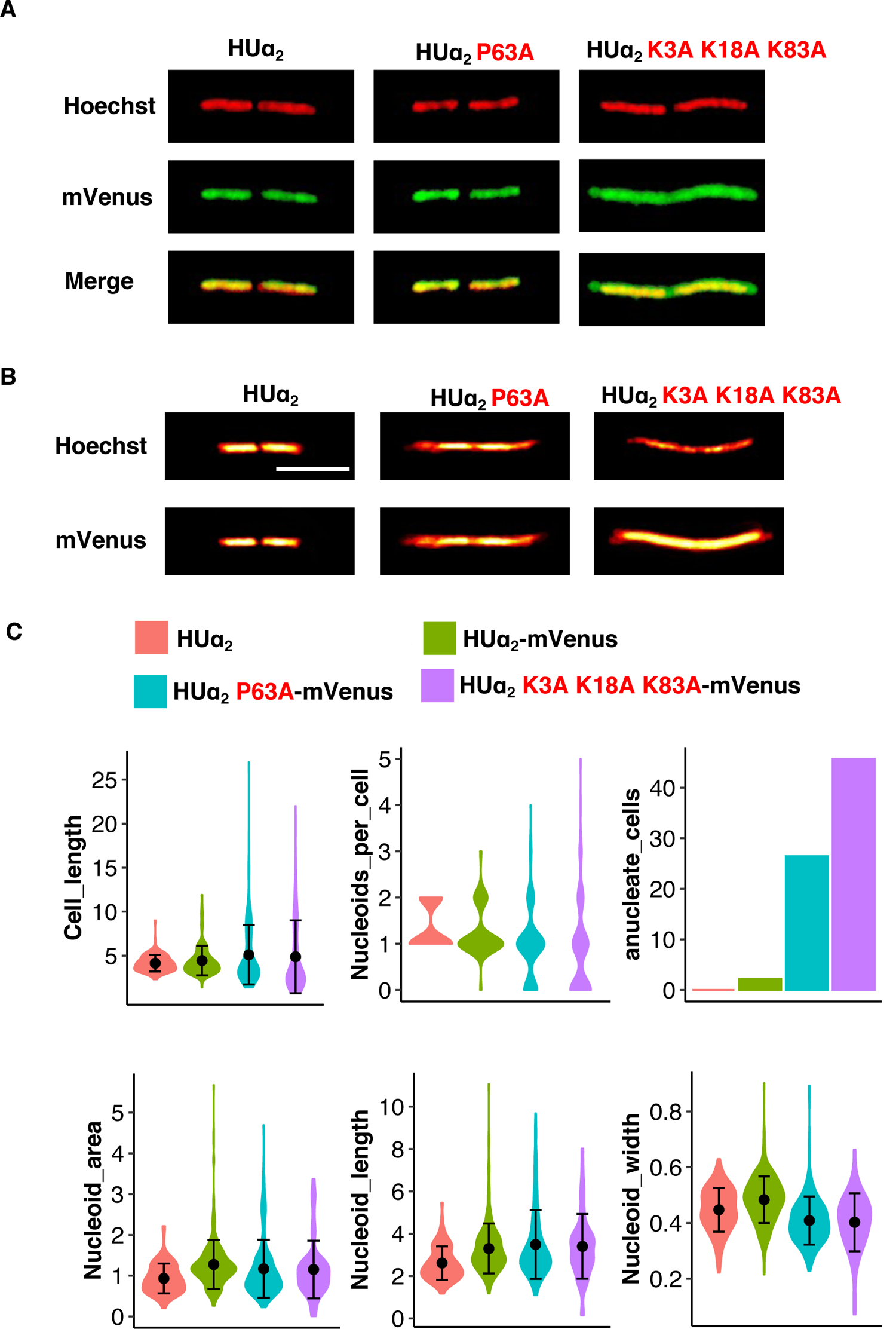
Nature of HU DNA binding in *E. coli* chromosome structure and function **A.** In vivo chromosome association of wild-type HUα_2_-mVenus or its variants. The fluorescence signal of mVenus is falsely colored as green. The DNA was stained with Hoechst 33342, and the fluorescence signal is falsely colored as red. The fluorescence images were obtained after subtracting the corresponding binary mask images from the original images. **B.** The aggregate images of the chromosome and the HUα_2_-mVenus or its variants, obtained by merging the fluorescence images of at least five individual cells with two or more well-segregated chromosomes. **(C)** Violin plots showing the distribution of cell length, the number of chromosomes per cell, chromosome area, width, and length of cells expressing HUα_2,_ HUα_2_-mVenus, or HUα_2_-mVenus variants. The bar graph shows the percentages of anucleate cells.

Second, we also investigated any influence of HU-binding on chromosome numbers per cell and chromosome morphology (width and length) along with associated cell length and cell morphology using fluorescence imaging, and if so, what kind of DNA binding is involved. Although we observed that the average cell length of the strain with HUα_2_ P63A was not much different from that of the strain with wild-type HUα_2,_ the distribution of the cell length varied significantly among individual cells (Figure 6C). The coefficient of variation in the mutant was about 66% compared to 38% in the wild-type. About 33% of the cells expressing HUα_2_P63A were longer than 5 µm, 17% of which were longer than 10 µm, compared to 24% of cells expressing wild-type HUα_2_ being longer than 5 µm with 6% of those being 10 µm longer.

Third, a more drastic effect of the P63A mutation occurred on the number of chromosomes per cell; about 26% of the mutant cells were anucleate (Figure 6C). The average length of the anucleate cells was 2.6 µm, about two times smaller than that of nucleated cells. Wild-type cells produced only 2% anucleate cells. We note that the anucleate cells are in small part due to the mVenus tag because the strain without tag produced no anucleate cells (Figure 6C). Nonetheless, the P63A mutation increased the fraction of anucleate cells by ten-fold compared to the wild-type HUα_2_ with the tag. The results suggest a role of HU structure-specific binding in chromosome partitioning. The triple lysine mutations K3A-K18A-K83A in HUα_2_ resulted in even higher fraction of anucleate cells (45%), presumably, adding to further loss of DNA-structure-specific binding.

After merging the images of individual cells containing at least two, or more, segregated chromosomes into a single aggregate image, the merged chromosome images showed separation between chromosomes in the strain expressing wild-type HUα_2_ (Figure 6B) suggesting that the chromosomes localize in the same position in each cell. The merged chromosomes, however, had no gap in between for the strain expressing HUα_2_ P63A or HUα_2_ with K3A-K18A-K83A mutations (Figure 6B), indicating that the disruption of HU-DNA interactions cause mis-localization of the chromosomes. Taken together, the production of anucleate cells and mis-localization of the chromosome in case of the P63A mutation alone suggest that the DNA structure-specific binding of HUα_2_ is critical for proper chromosome positioning and partitioning in *E. coli*. We did not observe a significant difference in average chromosomal area or average chromosomal length between the strains expressing wild-type HUα_2_ and mutant HUα_2_P63A (Figure 6C). The average chromosome width in the strain expressing HUα_2_ P63A, however, was significantly less than that of the strain with wild-type HUα_2_ (∼0.4 µm compared to ∼0.5 µm, p-value < 0.05, Figure 6C). These results show that the loss of structure-specific HU-DNA interactions are not only important for proper chromosome positioning and partitioning but also for maintaining the chromosome morphology.

## DISCUSSION

In *E. coli*, HU displays two kinds of DNA binding: (a) We have previously shown that the HU protein forms a crystal complex with a 20-bp DNA duplex of random sequence in which three highly conserved, surface-exposed lysine residues K3, K18, and K83 in the protein form ionic bonds with DNA phosphate (Hammel et al., 2016). In this report, we biochemically confirm the non-specific DNA binding of HUα_2_. (b) The physiologically relevant structure-specific DNA binding of HU was previously shown to require a proline residue P63 in the β-arm by crystal structure analysis (Guo and Adhya, 2007; Swinger et al., 2003). We also biochemically demonstrate that the structure-specific binding not only require the P63 but also involve the surface exposed lysine residues. We postulate that in the structurally specific binding of HUα_2_ to a distorted DNA, while the P63 residues in the tip of the β the non-specific contacts between the encompassing DNA and lysine residues on both sides of the bound HU dimer provides stability and sharper DNA bending (Figure 7).

**Figure 7.**
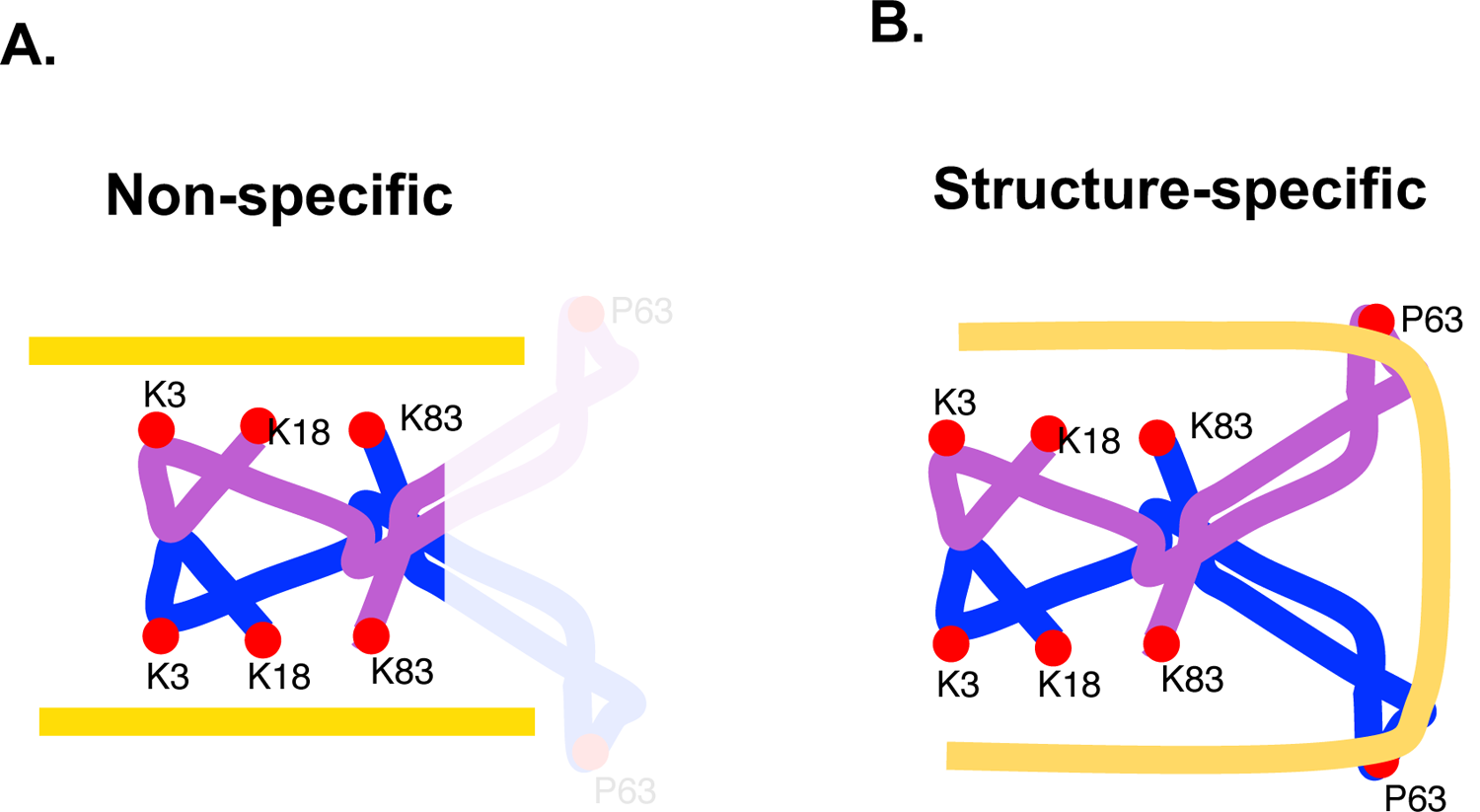
Cartoon representation of the non-specific and structure-specific DNA binding modes of HU. Two homo- or heteromeric subunits of HU are depicted as blue and purple. Critical amino acids for DNA binding are depicted as red circles. DNA is depicted in gold. In the non-specific binding mode to any B-DNA, three highly conserved lysine residues K3, K18, and K83 form electrostatic interactions with the phosphate backbone as shown previously by crystal structures and confirmed biochemically in this study. The bound DNA is mostly in the straight conformation. A single HU dimer can bind to two separate DNA molecules or two different segments of the same long DNA molecule. The beta arms do not contribute to the non-specific binding and therefore represented here in the faded color. In the structure-specific binding mode to DNA with distortions, the beta arms recognize a common structural motif of bent DNA helices wherein the proline residue P63 at the tip of the arms intercalates between bases at the site of kinks. As biochemically demonstrated in this study, the lysine residues contribute significantly to the binding. We propose that the non-specific interactions between the lysine residues and the phosphate backbone further along the DNA on both sides stabilize the binding and bends the DNA more sharply.

HU has a multifarious physiological role in *E. coli* involving DNA (Verma et al., 2019). Here we have assigned which type of HU binding to DNA – non-specific or structure-specific – is involved in many of its known physiological roles. Since HUα_2_ homodimer fulfils most of the roles of HUαβ heterodimer, we approached the objective by using HUα_2_ and its variants for technical ease by mutating the P63 and/or lysine residues in the *hupA* gene while deleting the *hupB* gene in the chromosome.

i. We have shown here that in restraining DNA supercoils, HU uses structure-specific binding, presumably by binding at any constituent cruciform structures.
ii. The deletion of HU has previously been shown to cause differential expression of many genes in *E. coli* (Prieto et al., 2012), but it is unclear how HU brings about the changes in gene expression - whether through HU-dependent chromosomal structural changes mediated by non-specific binding or by the formation of local higher-order DNA structures by structure-specific HU binding. We found that the DNA structure-specific binding, while not necessary for the overall chromosome structure, is critical for the regulation of gene expression by HU. We hypothesize that the structure-specific binding occurring at limited sites in the genome controls gene expression at transcription level by allowing a local DNA loop formation in the promoter region - repressosome or enhanceosome formation for gene repression or gene activation respectively (Geanacopoulos et al., 2001; Panne, 2008). For the assembly of repressosomes or enhanceosomes at specific genes, HU may recognize and bind with higher affinity to cruciform structures arising from inverted repeat sequences (IRS) in the negatively supercoiled *E. coli* genome (Sinden et al., 1980). The HU binding site (*hbs*) in the *gal* operon is an IRS, and negative supercoiling is required for repression of the *gal* operon (Lewis et al., 1999). We have confirmed here that the structure-specific HU binding is required for DNA loop formation leading to the *gal* operon repression. Since the P63 residue is also critical for the supercoil restraining ability of HU, the binding of HU to cruciform structures near genes may alternatively restrain negative supercoiling. There is indirect evidence suggesting that HU restrains transcription-induced supercoiling in the chromosome (Berger et al., 2016). The loss of HU binding can free these supercoils that, in turn, can alter transcription. Therefore, it is possible that drastic effect of the P63A mutation we observed on gene expression may have resulted from the formation of free supercoils. But the differentially expressed genes were not enriched for supercoiling sensitive genes identified previously (Peter et al., 2004) (Figure S4), contradicting the idea of supercoiling-mediated regulation of gene expression by HU.
iii. Although the structure-specific DNA binding of HU does not participate in the genome organization, as reported above, and discussed below, we have shown that HU-deleted strains are defective in proper chromosome partitioning giving rise to anucleate cells. The HU effect on chromosome partitioning is due to structure-specific binding as HUα_2_ with the P63 mutation showed chromosome partitioning defect. When studying the effect of HU in gene regulation, we observed that the genes encoding integration host factor (IHF) and DNA-binding protein from starved cells (Dps), plus the DNA gyrase inhibitor protein SbmC were down-regulated, whereas genes encoding MukB and the ParC subunt of DNA Topoisomerase IV were up-regulated upon the deletion of HU genes or by the disruption of the structure-specific binding in HUα_2_. MukB is the DNA binding subunit of the SMC complex MukBEF, which upon over-expression has recently been shown to form an axial core, resulting in the lengthwise compaction and organization of the *E. coli* chromosome (Makela and Sherratt, 2020a). TopoIV is a decatenase required for the segregation of the sister chromosomes. Additionally, the MukBEF complex and TopoIV physically interact, and the interaction likely coordinates timely segregation of the newly replicated chromosomes (Nolivos et al., 2016). Thus, we believe that the formation of anucleate cells and chromosome mis-positioning we observed due to the disruption of the structure-specific binding of HU may have resulted from the over expression of MukB and TopoIV. Here the HU structure-specific binding indirectly impacts the chromosome partitioning through alteration of gene expression.
iv. We found that only the non-specific binding participated in a major role of HU - its association with the chromosomal DNA that dictates the chromosome structure and organization. By tracking single molecule analysis in live cells, we have previously demonstrated (Bettridge et al., 2020) and further confirmed here that the non-specific binding of HU is responsible for this interaction with rapid association and dissociation kinetics that keep most HU within the nucleoid. Mutating the three lysine residues to alanine dramatically altered the interacting dynamics of HUα_2_ but mutating P63 to alanine did not have any effect (Bettridge et al., 2020). We do not rule out that structure-specific binding of HU at some defined sites may also play some architectural role in the chromosome as well.

In summary, our findings demonstrate that non-specific and structure-specific DNA binding modes of HU plays distinct roles: (a) HU has many targeted roles in various DNA metabolic processes like gene transcription, DNA supercoiling, chromosome positioning and partitioning. We have shown that some of these processes are direct, and some are indirect effect of structure-specific binding of HU to DNA. (b) The non-specific but highly dynamic binding of HU to chromosomal DNA organizes the chromosome that include wide-spread DNA-DNA contacts. The non-specific binding also keeps HU mostly as a component of the chromosome. HU is known to promote DNA-DNA contacts in the megabase pair range (Lioy et al., 2018; Walker et al., 2020) but the mechanism is unknown. We have previously shown that the non-specific DNA binding triggers multimerization of HUα_2_ (Hammel et al., 2016; Remesh et al., 2020). There is one HU dimer every ∼150 bp of the chromosomal DNA based on the estimated abundance of 30,000 HU dimers per cell during the exponential growth phase (Ali Azam et al., 1999), and most of the HU dimers are in the form of HUα_2_ (Claret and Rouviere-Yaniv, 1997). By consolidating these multiple lines of evidence, we argue that the multimerization of HUα_2_ molecules non-specifically bound to numerous positions on the chromosomal DNA brings distant genomic loci together. The rapid interacting dynamics of HU with DNA (Bettridge et al., 2020) may be necessary for the newly replicated genome to dynamically reorganize as the genome progressively segregate during DNA replication (Nielsen et al., 2006). Most target specific DNA binding proteins use base sequences specific motifs; their non-specific DNA binding abilities are relegated to as background noise. In contrast, the non-specific DNA binding activity of HU plays a critical role in chromosome architecture and organization, as also confirmed by recent our studies probing the 3D genome of *E. coli* using independent approaches (in preparation), making HU a unique DNA binding protein.

## ACKNOWLEDGEMENT

We thank Rupesh Kumar at Memorial Sloan Kettering Cancer Center for help in supercoiling assays.

## METHODS

### Key resource table

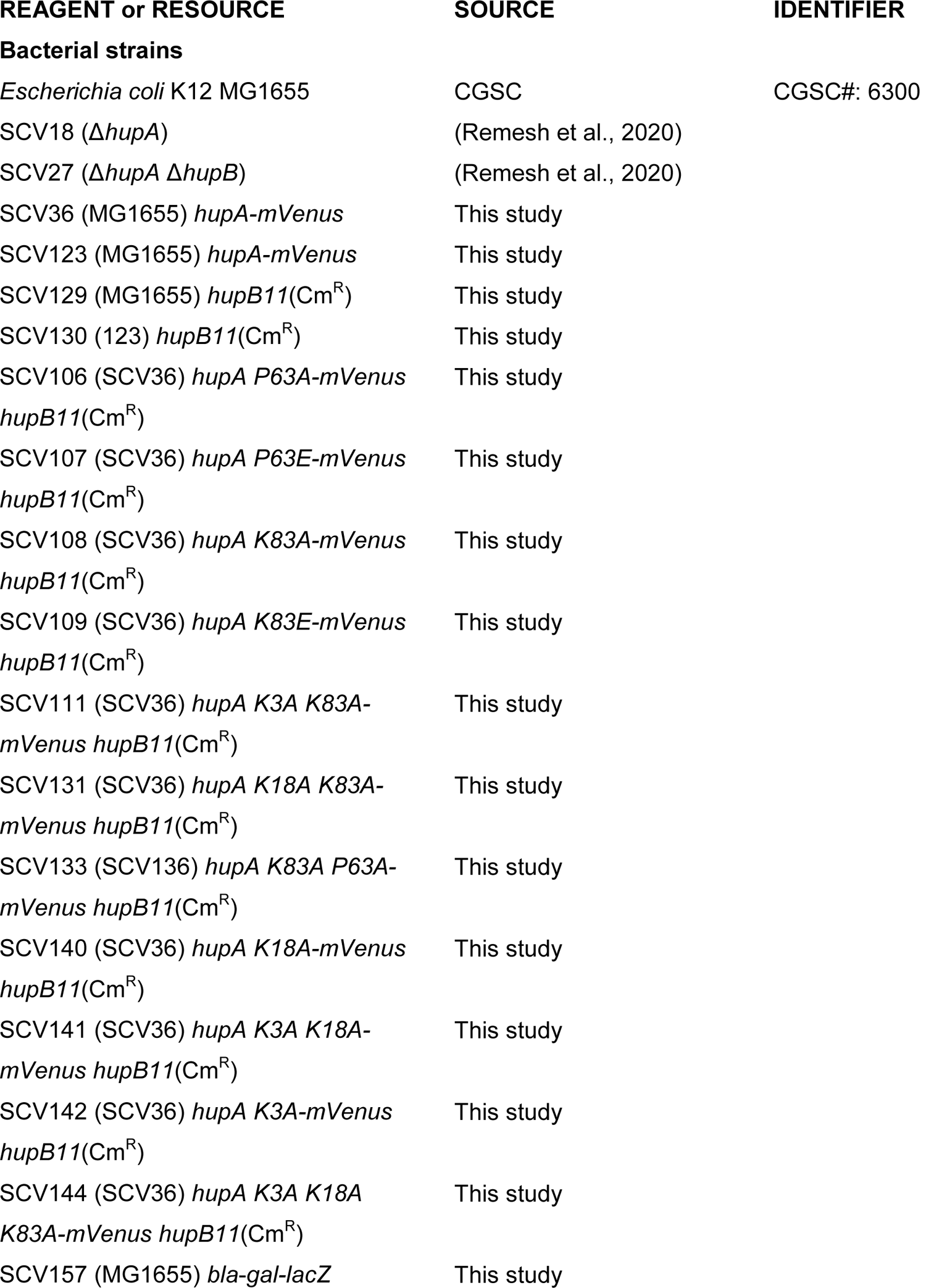
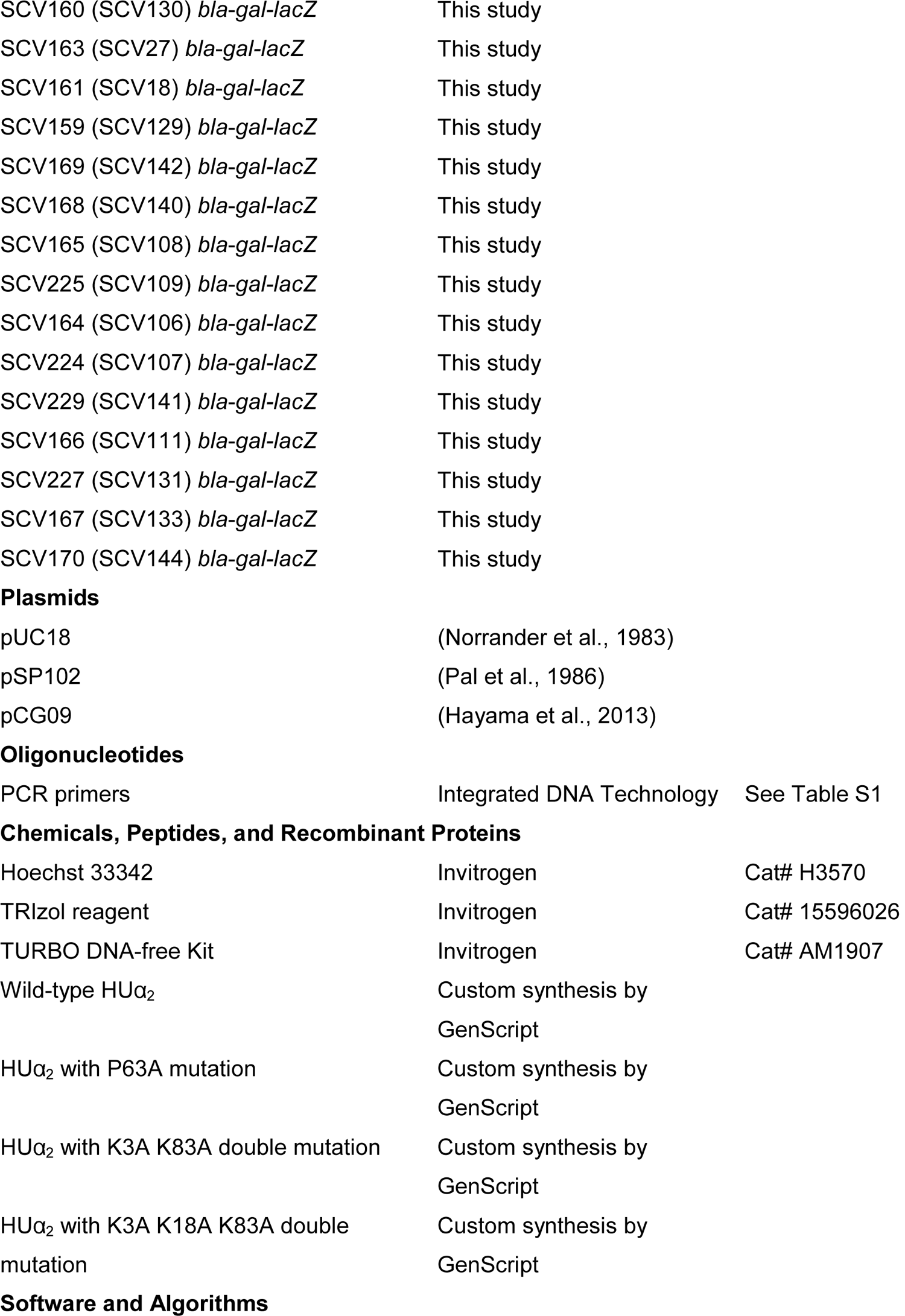
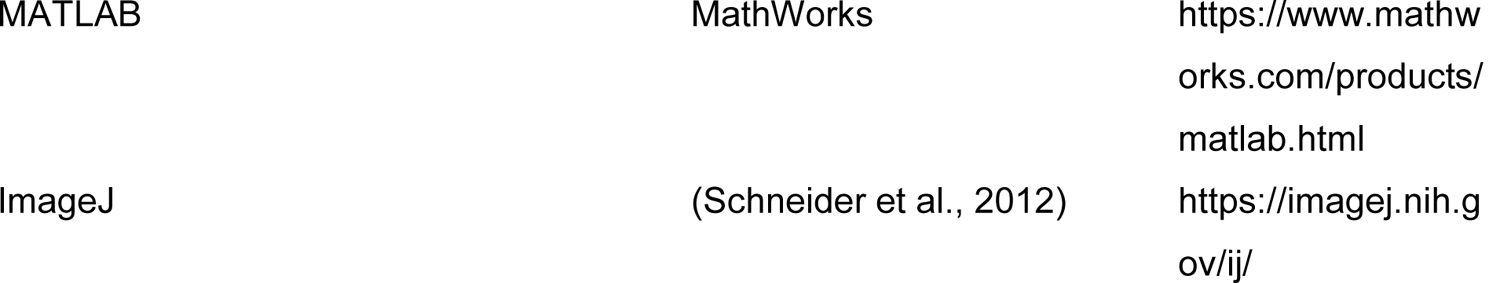

### Bacterial strains and growth conditions

Bacterial strains and primers are listed in Key Resources Table and Table S1, respectively. All strains were derivatives of *E. coli* K12 MG1655 (Blattner et al., 1997), and were constructed by Lambda red recombineering (Yu et al., 2000) using the plasmid pSIM6 (Datta et al., 2006). The SCV36 strain harboring *hupA-mVenus* fusion served as the parent strain for the strains harboring point mutations in the *hupA* gene. The *hupA-mVenus* fusion was constructed in two steps. In the first step, a pBAD-ccdB-kan genetic element that expresses CcdB toxin under the arabinose inducible promoter pBAD and encodes a kanamycin resistance gene was inserted after the stop codon of the *hupA* open reading frame. The recombinants were selected on LB agar plates containing 1% glucose and 30 µg ml^-1^ kanamycin. The recombinants were verified by PCR using the primers that bind outside the *hupA* gene as well as by the inability of the recombinants to grow on LB agar plates supplemented with 0.02% arabinose that induces the expression of the toxin CcdB. In the second step, a synthetic double-stranded DNA (synthesized by Integrated DNA Technologies) encoding the *hupA* gene, the glycine-serine-isoleucine (GSI) peptide linker, and yellow fluorescent protein mVenus was recombined. The recombinants were selected on LB agar with 0.02% arabinose and subsequently were verified by Sanger sequencing. After sequence verification, the plasmid pSIM6 was removed by repeatedly growing the strain at 37 C and selecting for ampicillin sensitivity. The point mutations were introduced using same steps; first inserting the pBAD-ccdB-kan element and then recombining the synthetic dsDNA encoding the *hupA*-mVenus fusion gene with desired mutations in the *hupA* gene. The *hupB11* allele (Wada et al., 1988) that has the *hupB* region between EcoRV and AatI sites replaced with a 1.4 kb HaeII DNA fragment containing the chloramphenicol-resistance (Cm^R^) gene of the pACYC184 plasmid was transferred from a strain of our lab collection using P1 phage transduction.

Cells were grown in M63 minimal media supplemented with 0.1% glycerol (v/v), 10 µg ml^-1^ thiamine, and 0.5 mg ml^-1^ casamino acids. For RNA extraction and fluorescence microscopy, cells were grown overnight, diluted 1000-fold and grown to A_600_ of 0.4-0.6.

### Protein purification

Wild-type HUα_2_ protein and HUα_2_ protein with P63A mutation, K3A, K83 double mutations, or K3A, K18A, KA83 triple mutations were purified by GenScript Biotech Corporation. 6x-histidine tag containing Enterokinase cleavage site was placed at the N-terminus. Proteins were expressed in *E. coli* strain BL21(DE3) and purified with >95% purity using Ni-affinity column followed by Q-Sepharose and size-exclusion chromatography. The proteins were stored in the buffer containing 50 mM Tris HCl pH 8.0, 150 mM NaCl, 10% Glycerol. The tag was removed by histidine tagged Bovine Enterokinase (GenScript Biotech Corporation) before using the proteins for DNA binding assays.

### DNA binding assays

The 6-carboxy fluorescein (6-FAM) labelled duplex DNA synthesized by Integrated DNA Technologies were used for DNA binding. The following oligonucleotides were used for synthesizing the duplex DNA.

B-form DNA of random sequence: CCGACTAAGTACATGTGAGAATTTTGCTGCCTTCGAACCT and /56-FAM/AGGTTCGAAGGCAGCAAAATTCTCACATGTACTTAGTCGG

Cruciform DNA: CCTAGCAAGGGGCTGCTACCTTTGGTAGCAGCCTGAGCGGTGG and /56-FAM/CCACCGCTCAACTCAACTGCTTTGCAGTTGAGTCCTTGCTAGG

The binding assays were carried out in the 100 µl reaction volume containing 50 mM Tris-HCl pH 8.0, 30 mM NaCl, 1 nM DNA, and varying concentrations of the protein. The reactions were incubated in 6 mm x 50 mm glass test tubes at room temperature for 10 min and then fluorescence polarization measurements were taken in the Beacon™ 2000 instrument at 25℃ Subsequently, 20 µl reaction volume was loaded onto 6% DNA retardation gels (Invitrogen), and electrophoresis was carried out at room temperature in 0.5x Tris borate buffer, pH 8.0. The gels were imaged using Bio-Rad ChemiDoc MP Imaging System. The milipolarization (mP) units were plotted in GraphPad Prism v9. The data points were fitted using the Hill slope equation 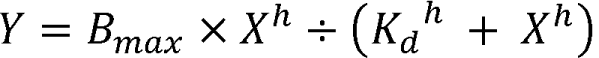 wherein B_max_ is the maximum binding in the mP units, *X* is the protein concentration, *h* is the hill slope, and *K_d_* is the protein concentration needed to achieve the half-maximum binding at equilibrium.

### Transcription Assays

In vitro transcription reactions were carried out as described earlier (Lewis, 2003). Supercoiled plasmid pSA850 (40 nM) was preincubated at 37°C for 5 min with 20 nM RNA polymerase and 200 nM GalR and/or varying concentrations of wild-type HUα_2_ or its mutant variants in a total μl containing transcription buffer (20 mM Tris acetate/10 mM Mg acetate/50 mM NaCl) supplemented with 1 mM DTT, 1 mM ATP, and 0.8 units recombinant ribonuclease inhibitor (rRNasin. As needed, were added before RNA polymerase. Transcription reactions were initiated by adding nucleotides to a final concentration of 0.1 mM GTP and CTP, 0.01 mM UTP, and 5 μCi [α 32P] UTP (1 Ci = 37 GBq). The reactions were incubated for an additional 10 min before they were terminated by the addition of an equal volume (50 μ loading dye (90% formamide/10 mM EDTA/0.1% xylene cyanol/0.1% bromophenol blue). Samples were heated to 90°C for 2–3 min, chilled, then loaded on an 8% sequencing gel and electrophoresed at a constant power of 60 W in TBE (90 mM Tris/64.6 mM boric acid/2.5 mM EDTA, pH 8.3). The RNAI transcripts (106 and 108 nts) were used as an internal control to normalize the relative amount of transcript from *P2* promoter.

### Supercoiling assay

The singly nicked pCG09 plasmid was prepared by incubating CsCl purified negatively supercoiled pCG09 (60 μ l of 1X NEB Smart Cut buffer with the nicking endonuclease Nb.BbVCI (NEB, 8.5 units) at 37 °C for 30 min. The plasmid was purified by phenol: chloroform: isoamyl alcohol extraction, recovered by standard ethanol precipitation, and dissolved in 1X buffer containing 10 mM Tris-HCl (pH 8.0), 1 mM EDTA, and 0.05 % (v/v) IGEPAL 630 (Sigma). Wild-type HUα_2_ protein and its variants were incubated with singly nicked pCG09 plasmid for 5 min in a buffer containing 50 mM HEPES-KOH (pH 7.5), 20 mM KCl, 30 mM NaCl (from protein stocks containing 150 mM NaCl), 10 mM DTT, 5 mM MgOAc, 7.5 % glycerol, 2 mM ATP-Mg and 5 ng/µL tRNA. T4 DNA ligase was added to seal the nick. The reactions were stopped by addition of NaCl and EDTA followed by deproteination with SDS and proteinase K for 30 min at 37℃. Topoisomers were resolved on a 0.8 % agarose gel at room temperature and 23 volt for 18 hours. Gels stained with SYBR Gold and imaged by Typhoon scanner.

### β-galactosidase assay

The *gal-lacZ* reporter was constructed by replacing the chromosomal region of MG1655 starting from the *lacI* up to the start codon of the *lacZ* with the DNA fragment of the plasmid pSA813 containing ampicillin resistance gene *bla*, the *gal* promoter, and the *galE* gene. The *gal-lacZ* reporter was transferred to other strains using P1 phage transduction.

β-galactosidase assay was carried out using standard protocol. Briefly, cells were diluted in Z buffer and permeabilized by adding 100 μl chloroform and 50 μ 0.1% sodium dodecyl sulfate. The reaction was initiated by adding 0.2 ml of 4 mg/ml o-nitrophenyl-β-D-galactoside in 0.1M phosphate buffer (pH 7.0), incubated at 28°C, and stopped after sufficient yellow color developed by adding 0.5 ml 1M Na_2_CO_3_. 1 ml of the reaction was transferred to an eppendorf tube, spun at maximum speed to remove debris and chloroform, and the absorbance at 420 nm was recorded for each tube. The following equation was used to calculate units of enzyme activity: 1000 x (A_420_) / (A_600_ x T x V) where T is the time of reaction in minutes and V is the volume of cells used in the reaction in milliliters.

### RNA sequencing

For total RNA isolation, a frozen pellet of cells was resuspended in 1 ml TRIzol reagent (Life Technologies) to homogenization and incubated at room temperature for 5 min. To the resuspension, 0.2 ml chloroform was added and mixed by inverting the tube for 15 seconds. The mixture was incubated at room temperature for 10 min and then centrifuged at 20,000 x g for 10 min at 4 °C. After centrifugation, ∼0.6ml of the upper phase was transferred to a new centrifuge tube containing 0.5 ml isopropanol. The mixture was incubated at room temperature for 10 min and then centrifuged at 20,000 × g for 15 min at 4 °C. After centrifugation, the supernatant was discarded and the pellet was washed twice with 1ml 75% ethanol, made with Diethyl pyrocarbonate (DEPC)-treated water, by centrifugation at 13,544 x g, for 5 min at 4 °C. After the second wash, the tube was left open for 10–15 min at room temperature to dry the pellet. To the pellet, 50 the tube was left at 37 °C for 10–15 min and then the pellet was fully resuspended using a pipette. DNA was removed using TURBO DNA-free Kit (Invitrogen). Quality of total RNA was determined by electrophoresis on the TapeStation system (Agilent). Paired-end sequencing libraries were prepared with 2.5 g of total RNA using Illumina TruSeq Stranded Total RNA library prep workflow with Ribo-Zero. Samples were pooled and sequenced on HiSeq4000 with read length of 150. Samples were barcode demultiplexed allowing one mismatch using Bcl2fastq v2.17. The reads were trimmed for adapters and low-quality bases using Cutadapt software. Alignment of the reads to the annotated transcriptome of *E. coli* K12 MG1655 was done using STAR. Transcript abundances were calculated by RSEM, and differential expression analysis was done using the glmTreat function of edgeR. We identified differential expression based on false discovery rate (FDR) cut off of 0.05 and log_2_ (log with base 2) fold change of 1.5. The RNA-Seq data have been deposited to the Gene Expression Omnibus (GEO) data base and can be accessed with the GEO accession number GSE175465.

### Mu phage growth assay

Strains were first grown overnight in LB broth. The overnight cultures were 100 times diluted into LB broth supplemented with 1mM CalCl_2_ and 2.5 mM MgCl_2_ and grown to A_600_ between 0.3-0.4. Cells were centrifuged and resuspended such that A_600_ is 1.0. 100 µl cells were mixed with 10 µl serially diluted phage lysate and incubated at 37℃ for 15 min. The mixture was added to 3 ml of 0.7% top agar and poured onto Terrific Broth agar plates. The next day, plaque were counted by eye. The plaque forming units/ml was calculated as average number of plaques divided by dilution of the phage lysate times volume of the phage lysate.

### P1 plasmid transformation assay

Electrocompetent cells prepared using standard protocol were transformed with pUC18 or pSP102 plasmid DNA by electroporation. The transformants were selected on LB agar plates containing appropriate antibiotic. The transformation efficiency was calculated as colony forming units per microgram plasmid DNA.

### Fluorescence microscopy

1 ml bacteria culture was stained with 10 µg ml^-1^ Hoechst 33342 for 10 min and centrifuged at 1000 x g for 3 min at room temperature. All supernatant was removed except the volume in microliters equal to A_600_ x 200. The resuspended cells were stained with 15 µg ml^-1^ FM464 for 10 min. Cells were then spotted onto 35 mm Poly-D-Lysine coated glass petri dish (MatTek Life Sciences) and covered by M63 glycerol 1% (w/v) agarose pad. Z-sections were collected using DeltaVision imaging system (GE Healthcare) equipped with CoolSNAP_HQ2 camera. The pixel size of each image was 0.064 µm x 0.064 µm x 0.2 µm. mVenus fluorescence was detected using Fluorescein isothiocyanate (FITC) filter (excitation: 475/28; emission: 525/48) with 100% light transmission. Hoechst 33342 was detected using 4′ diamidino-2-phenylindole (DAPI) filter (excitation: 390/18; emission: 435/48) with 50% light transmission. FM464 was detected using Tetramethylrhodamine filter (excitation: 542/27; emission: 597/45) with 100% light transmission. Exposure time for Hoechst 33342 and FM464 was 0.5 s, and for mVenus was 0.2 s. The images were deconvoluted using the recommended method in the SoftWoRx software. Image quantitation was performed in ImageJ using the middle section of the z-stack. Cell length was manually measured using FM464 fluorescence. Chromosome length and chromosome width were measured as major axis and minor axis, respectively.

To generate the aggregate images of the chromosome and the HUα_2_-mVenus, the images of individual cells were cropped and rotated such that the long axis of cells corresponded to the x-axis and the middle of the cell corresponded to the axis origin. Then, the chromosome and HUα_2_-mVenus channels were split, channel images of individual cells were concatenated, and aggregated into a single image using 2-D sum intensity projection. The aggregate images were cropped to 12.99 micron long and 4.67 micron wide. For better visual comparison, the pixel intensity of the aggregate images was normalized in MATLAB by the maximum pixel intensity of that image, and then multiplied by 255. To account for intensity and noise differences, the aggregate images were further normalized in MATLAB using the aggregate image of WT chromosome as template and using histogram matching combined with gradient information (Sintorn et al., 2010).

---

**Figure S1.**
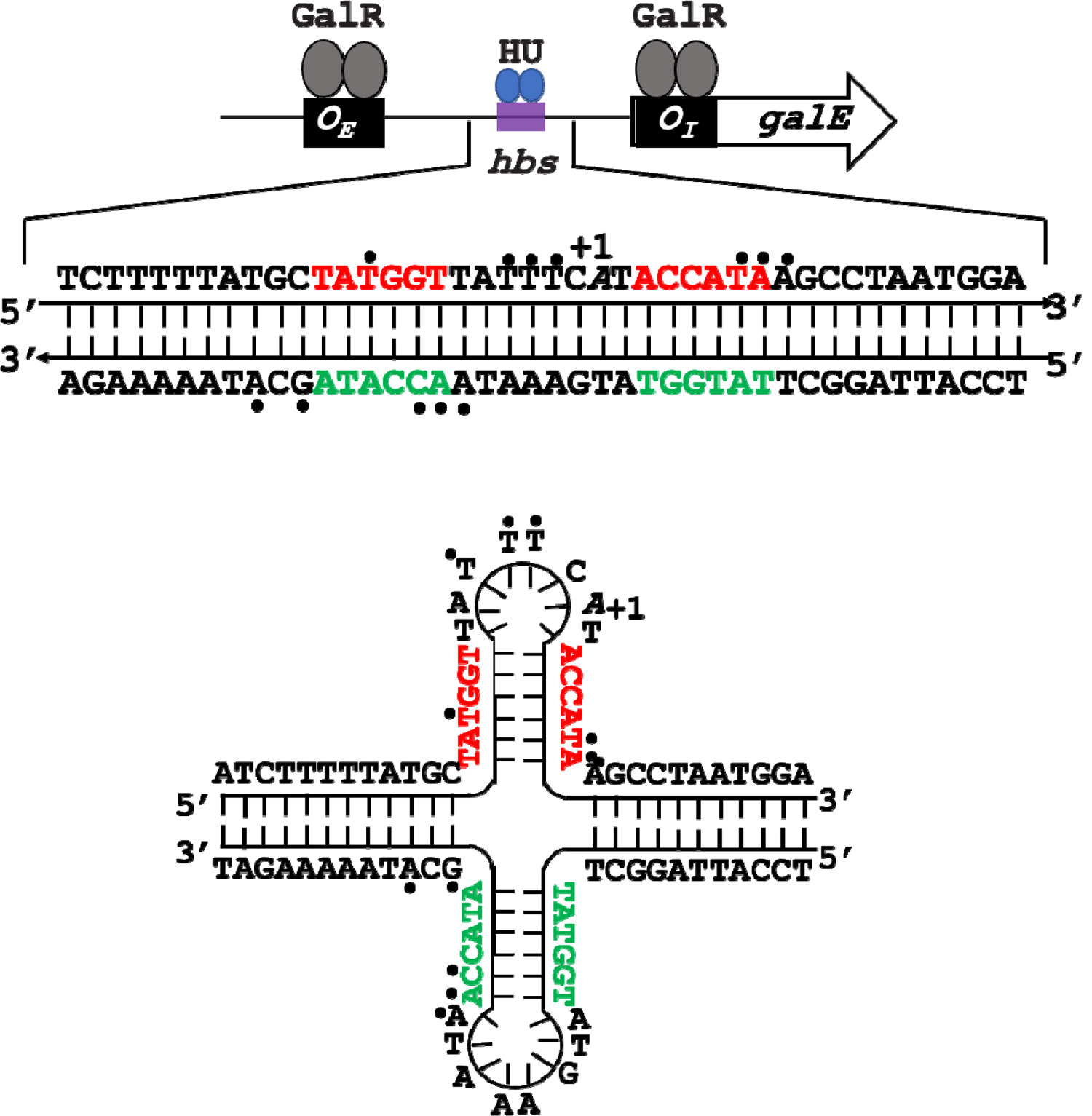
The presence of a potential cruciform structure within the HU binding site (*hbs*) at the *gal* operon. The inverted repeat sequence is highlighted in red and green. Black dots represent nucleotides in physical contact with HU in a supercoiled DNA template as determined previously by chemically converting HU into a nuclease (Aki and Adhya, 1997). +1 is the transcription start site.

**Figure S2.**
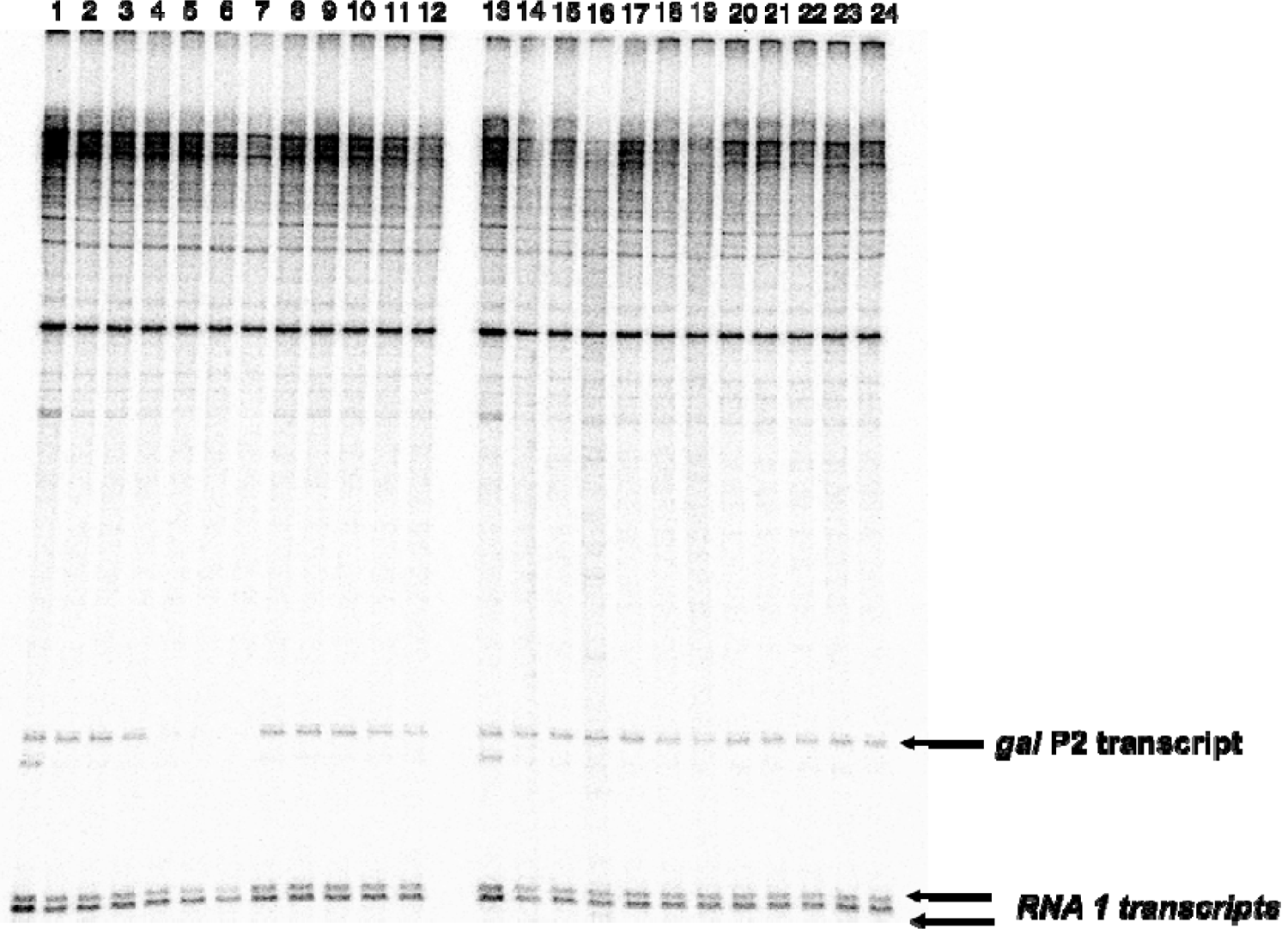
Effect of point mutations in HU on the in vitro transcription from the *gal* promoters. Lanes 1 and 13: no GalR or HU; Lanes 2 and 14: 200 nM GalR, Lanes 3 to 7: 200 GalR and 50, 100, 200, 400, and 800 nM wild-type HUα_2_; Lanes 8 to 12: 200 GalR and 50, 100, 200, 400, and 800 nM HUα_2_ P63A; Lanes 15 to 19: 200 GalR and 50, 100, 200, 400, and 800 nM HUα_2_ K3A K83A; Lanes 20 to 24: 200 GalR and 50, 100, 200, 400, and 800 nM HUα_2_ K3A K18A K83A.

**Figure S3.**
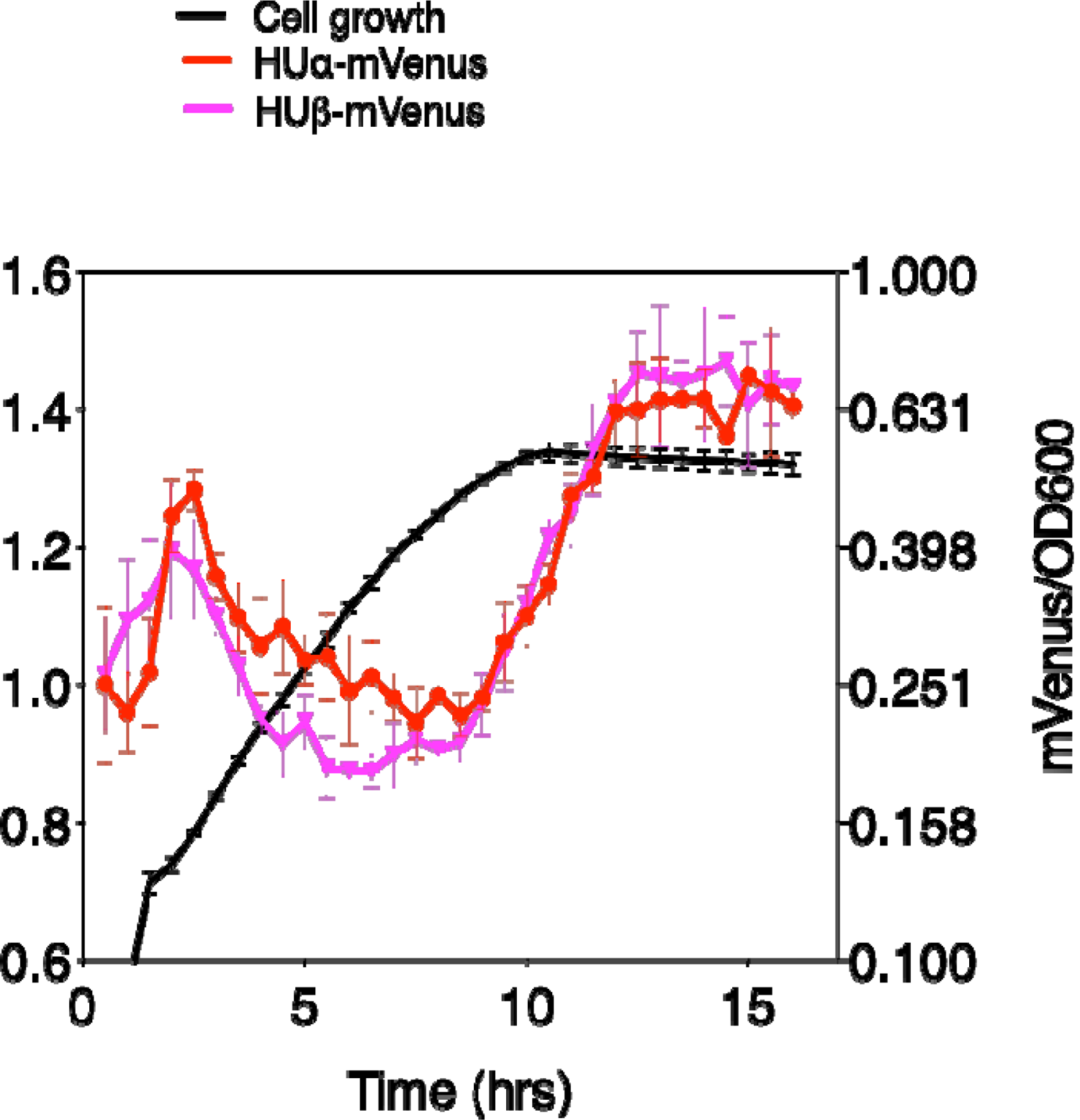
Levels of α and β subunits of HU during cell growth Strains expressing HUα-mVenus or HUβ-mVenus were first grown in standard test tubes in M63 minimal m^α^edia supplemen^β^ted with 0.1% glycerol (v/v), 10 µg ml^-1^ thiamine, and 0.5 mg ml^-1^ casamino acids. Cultures were diluted 100-fold into fresh media, and 1-ml added to 24-well plates with glass bottom (Greiner Bio-One). Cell growth and fluorescence were measured in Tecan Spark plate reader in real time every 30 min. Please note that optical density at 600 nm measured by the plate reader is not the same as that measured by standard spectrophotometer. Since HUα and HUβ have a higher affinity with each other than between themselves (Ramstein^α^et al., 20^β^03), WT strain likely produces HU predominantly in the form of the HUαβ heterodimer throughout the growth cycle.

**Figure S4.**
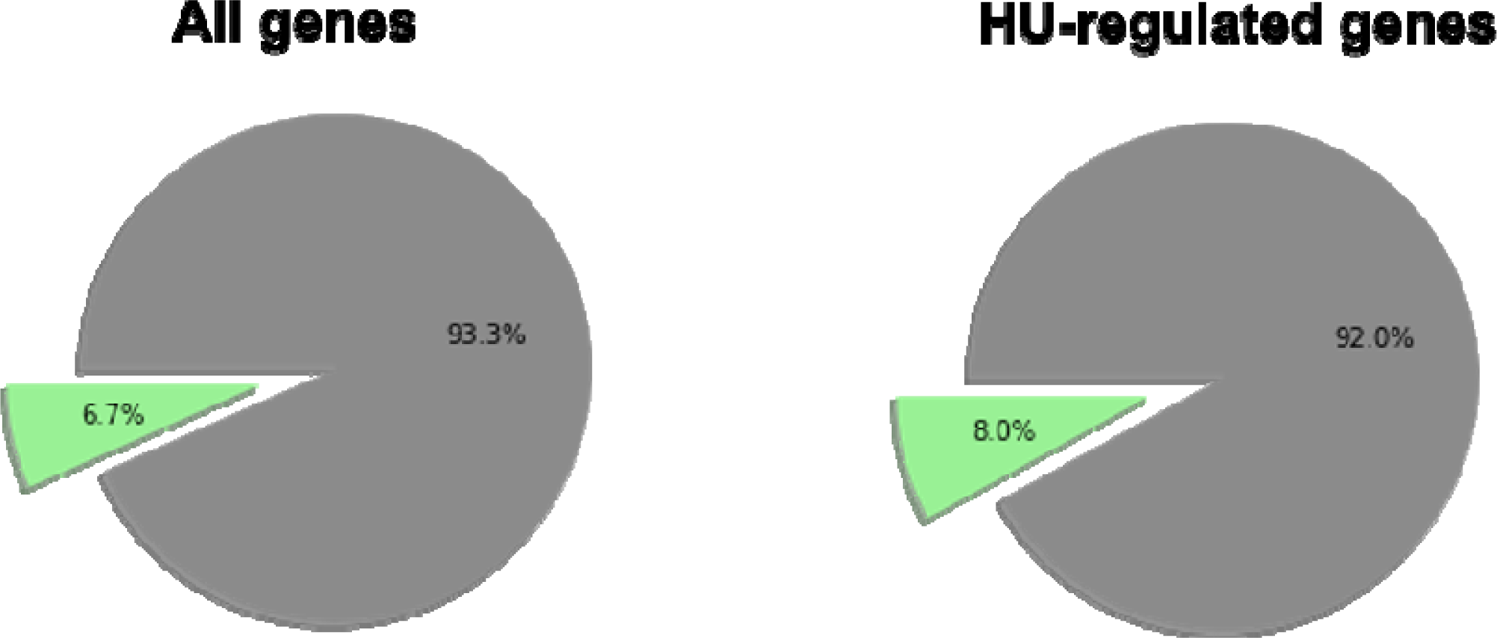
Supercoiling sensitive genes. Pie chart showing supercoiling sensitive genes ((Peter et al., 2004) as percentage of total *E. coli* K12 MG1655 genes or differentially expressed genes identified in the absence of HU.

**Table S2.**
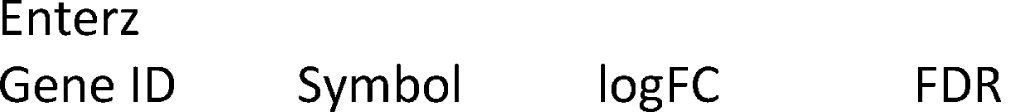
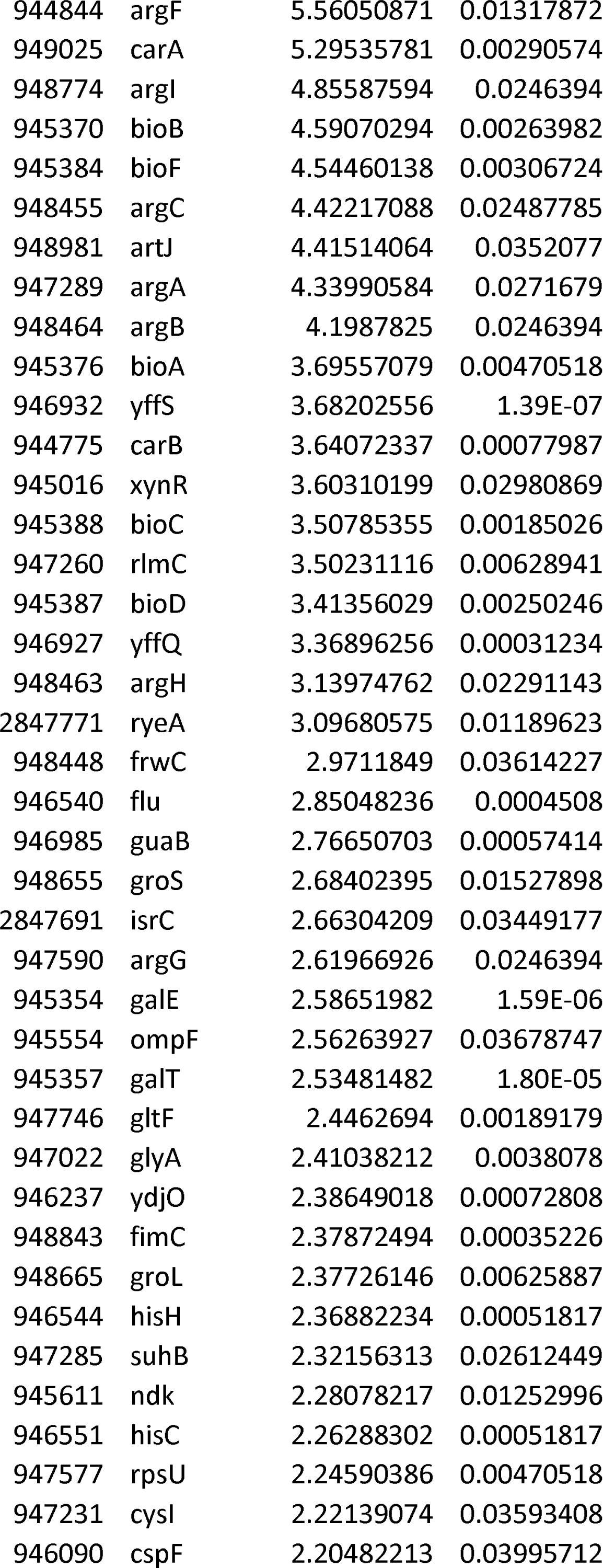
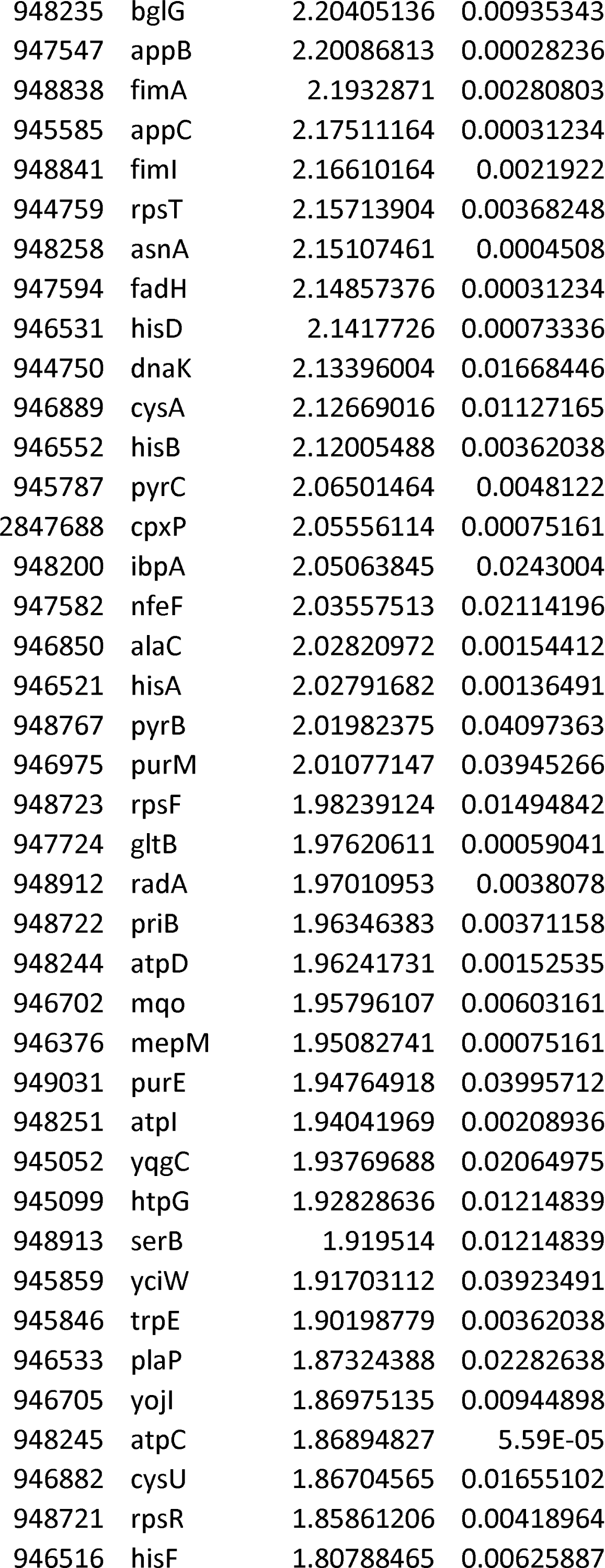
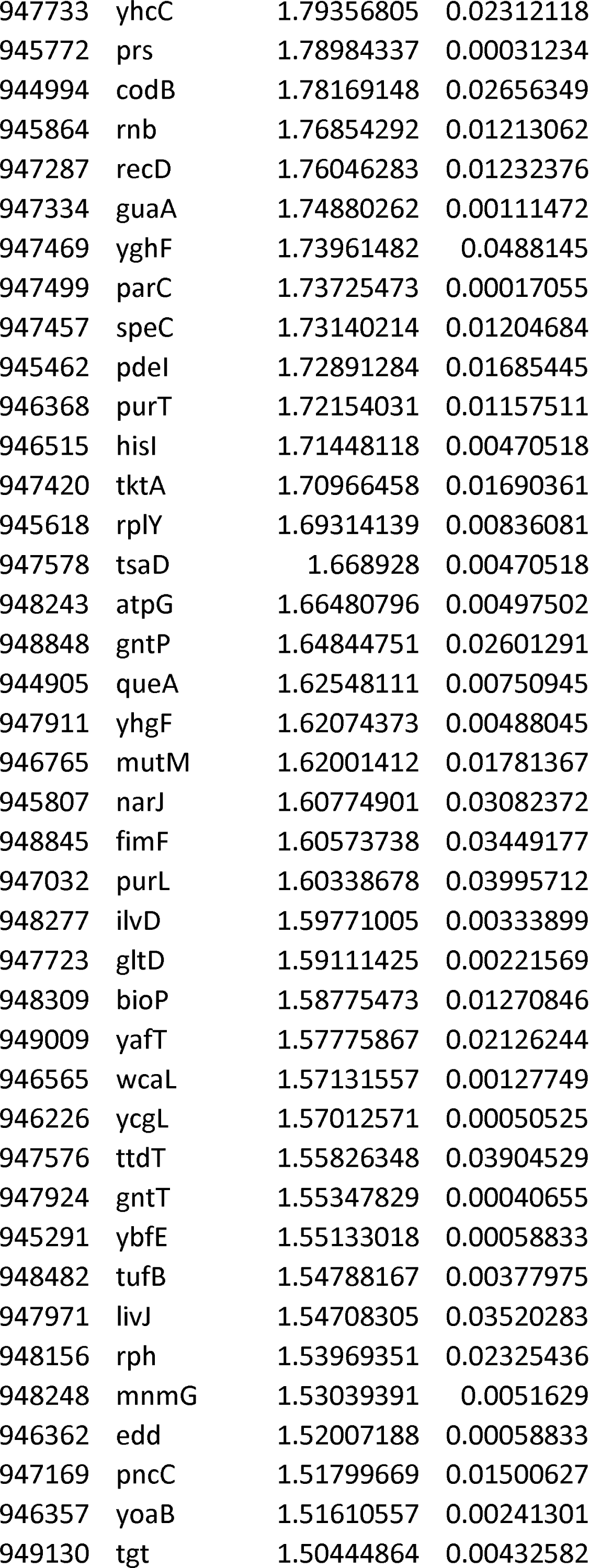
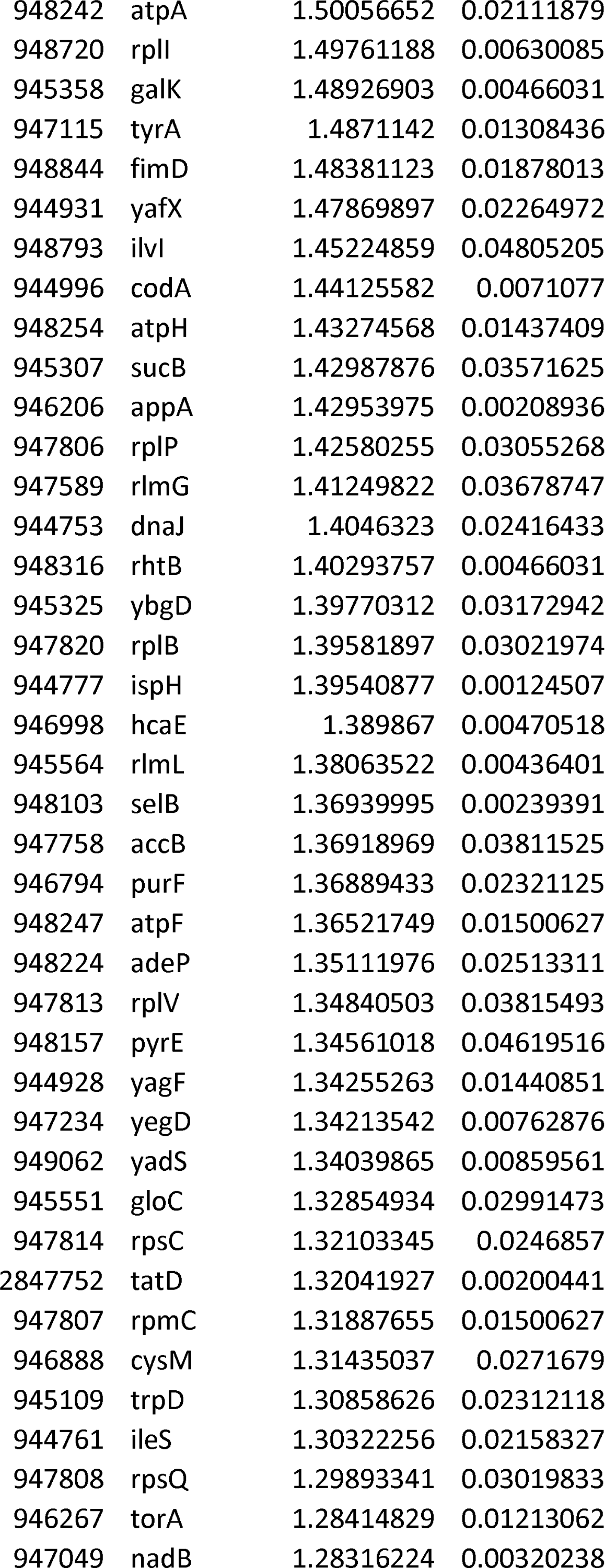
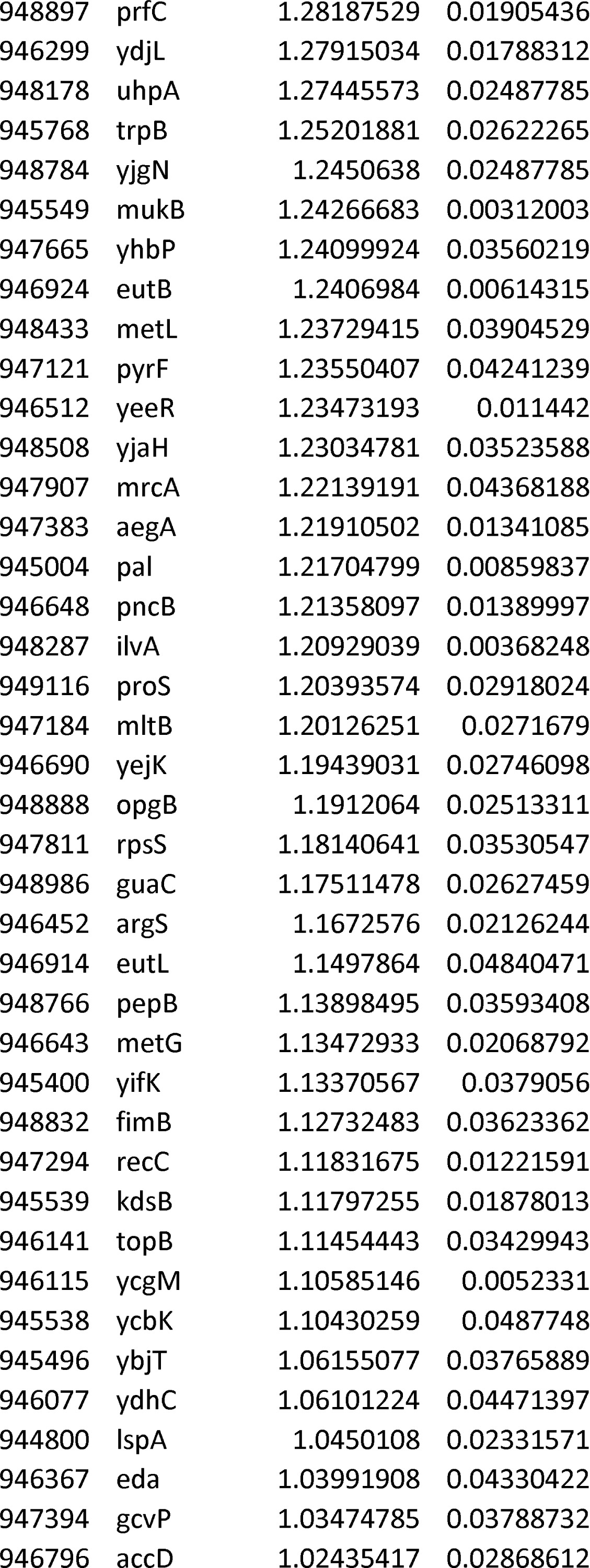
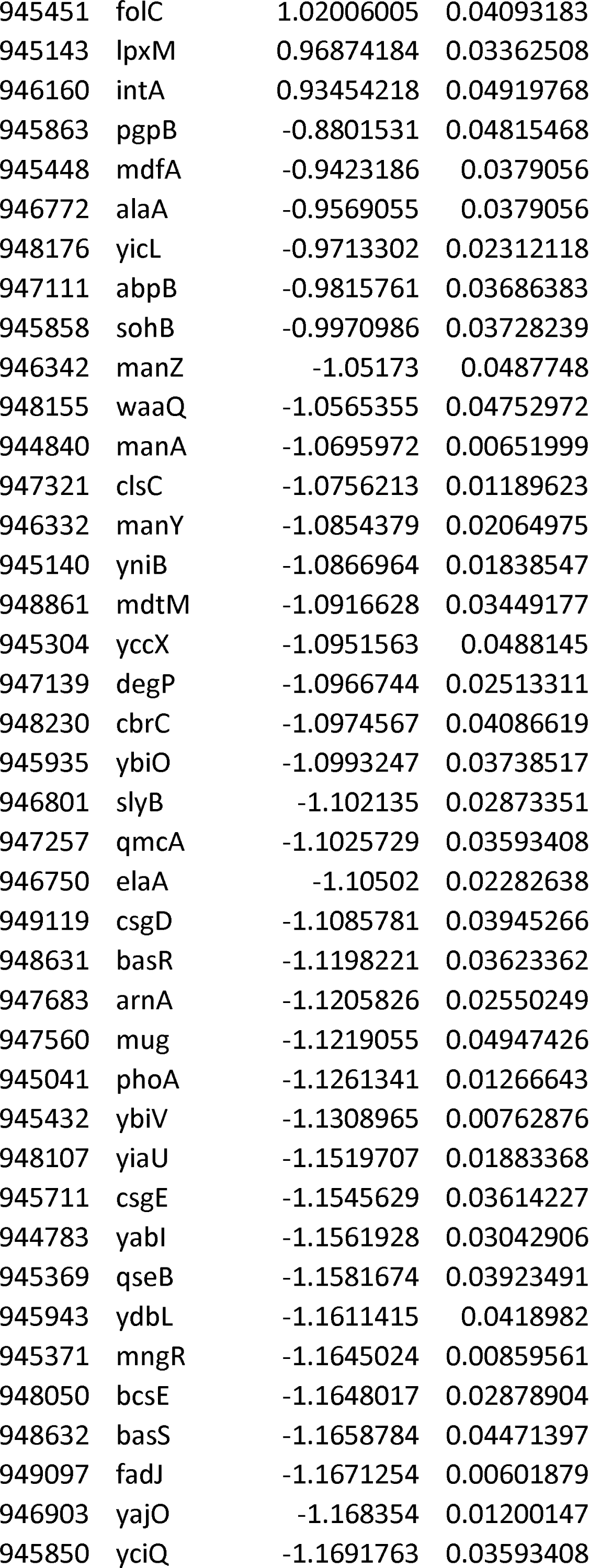
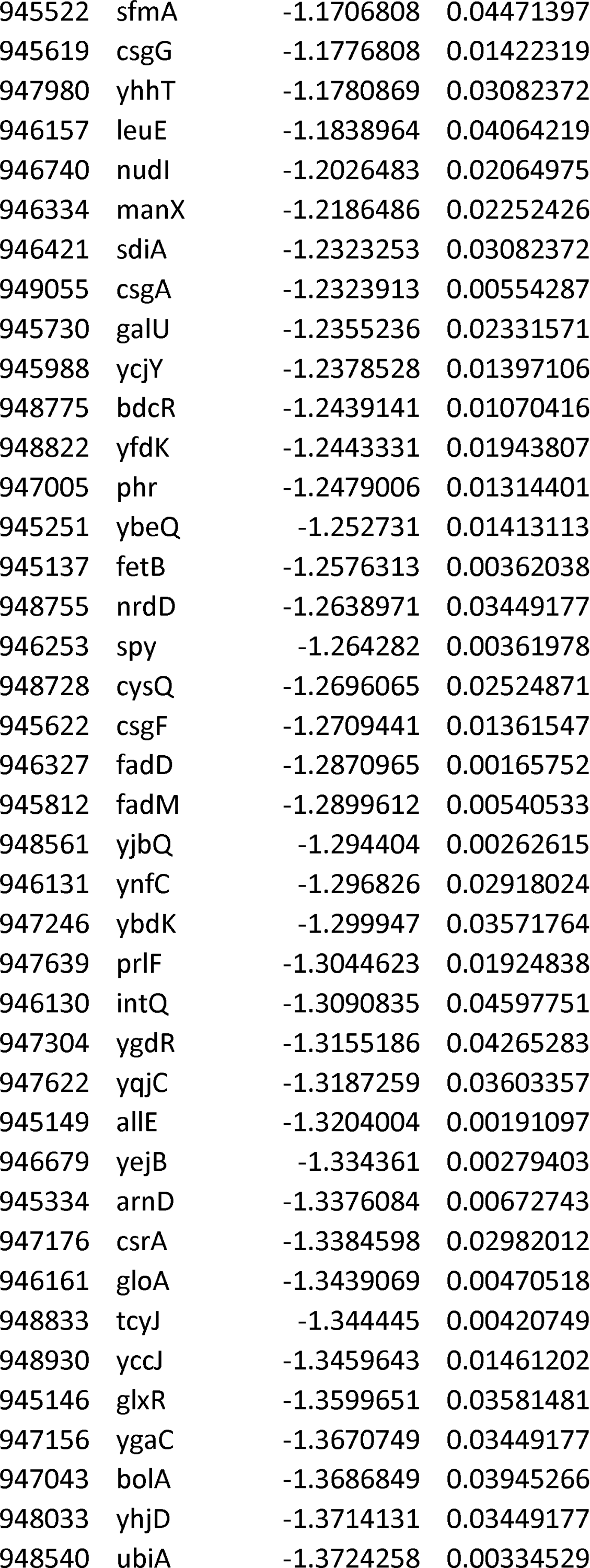
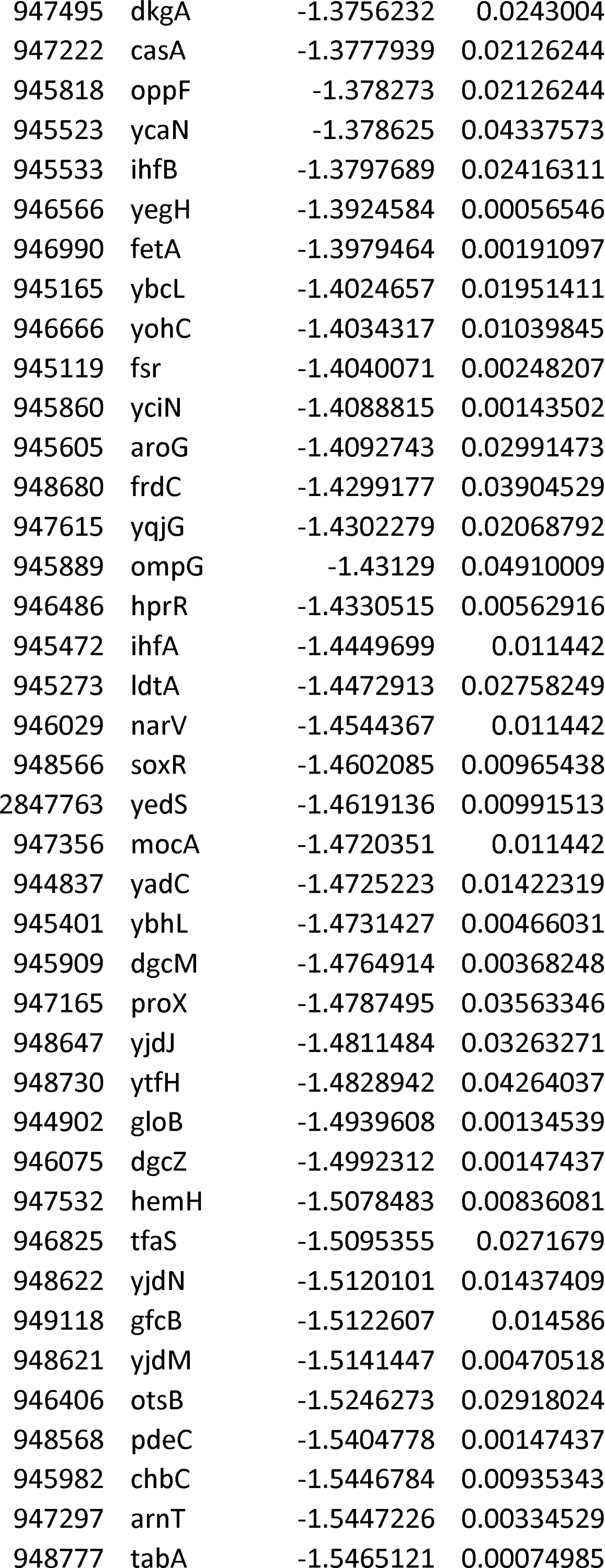
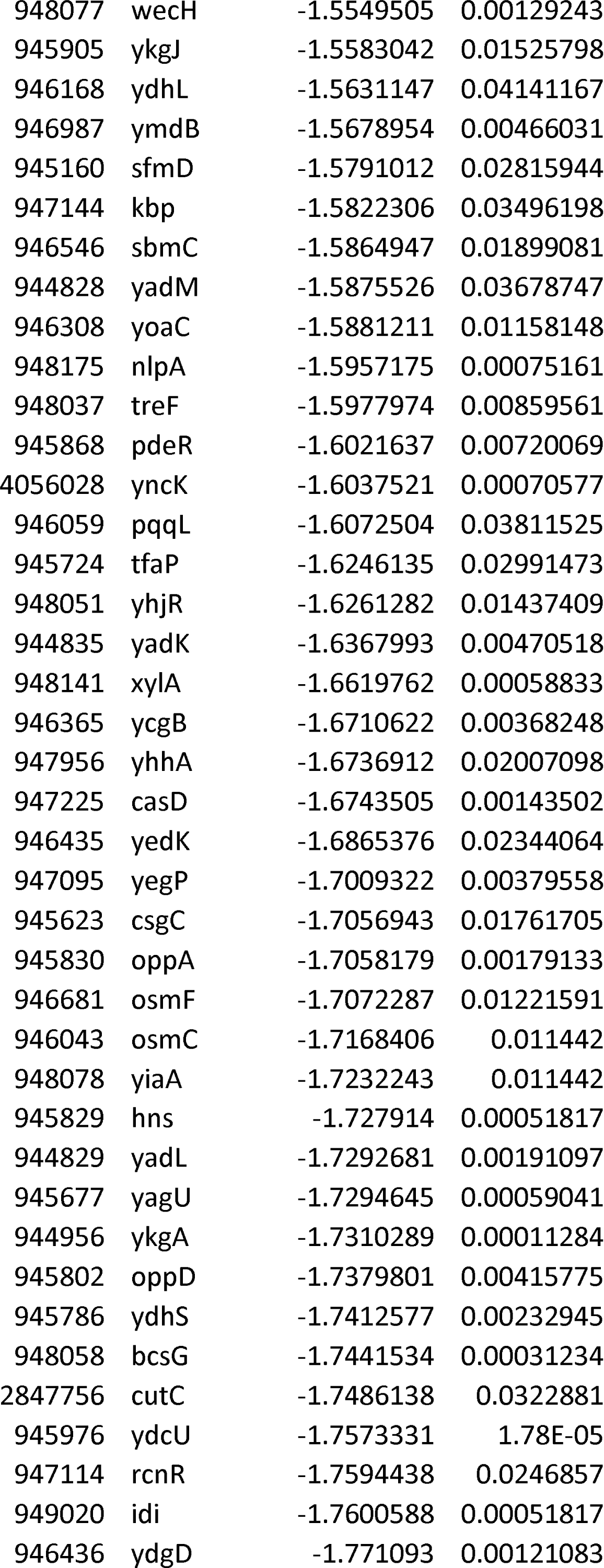
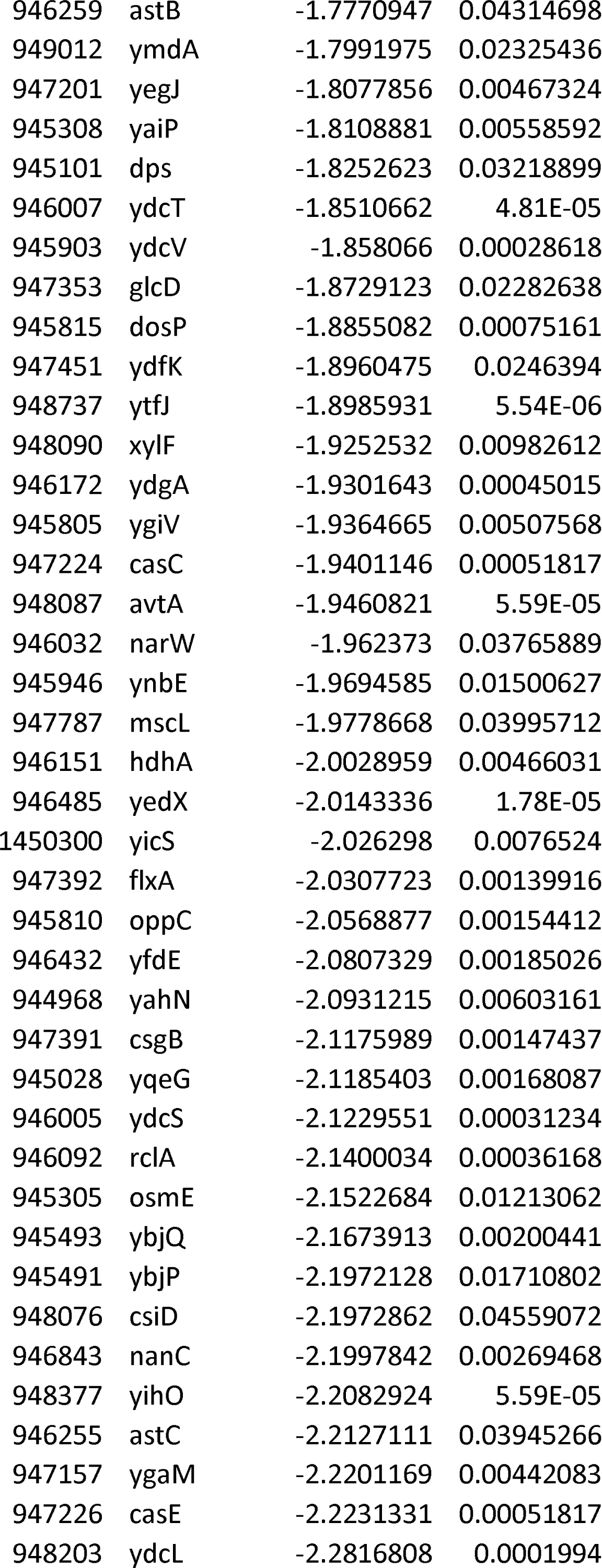
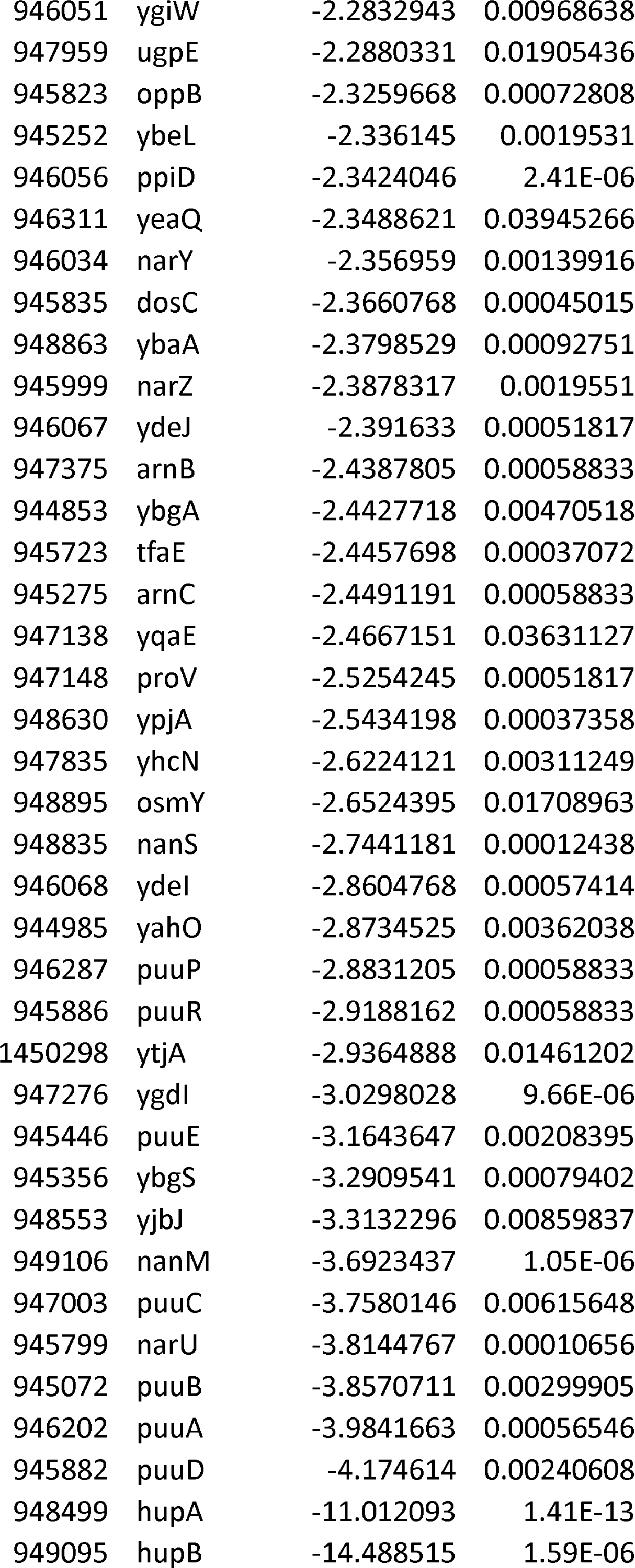
Differentially expressed genes in the HU null strain relative to strain expressing wild type *hup* genes.

**Table S3.**
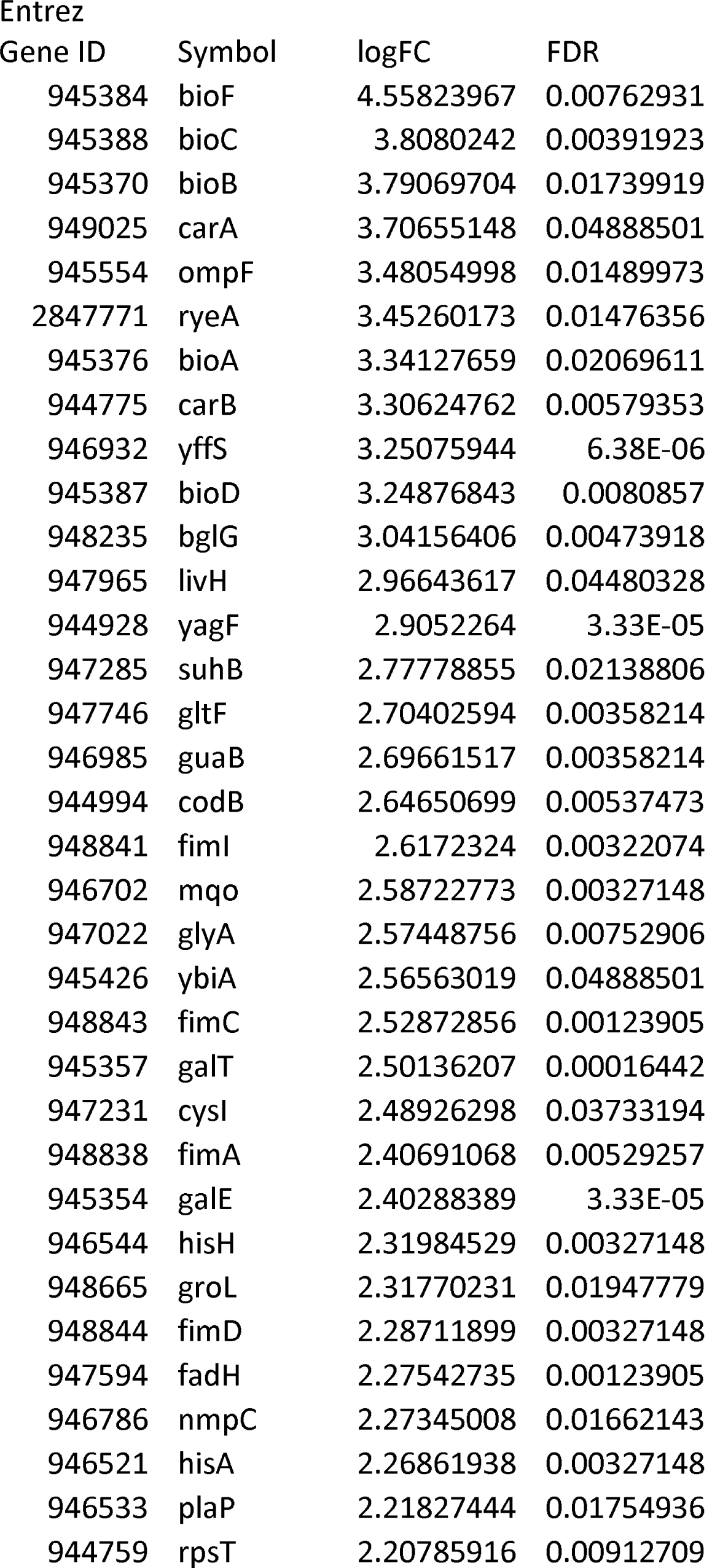
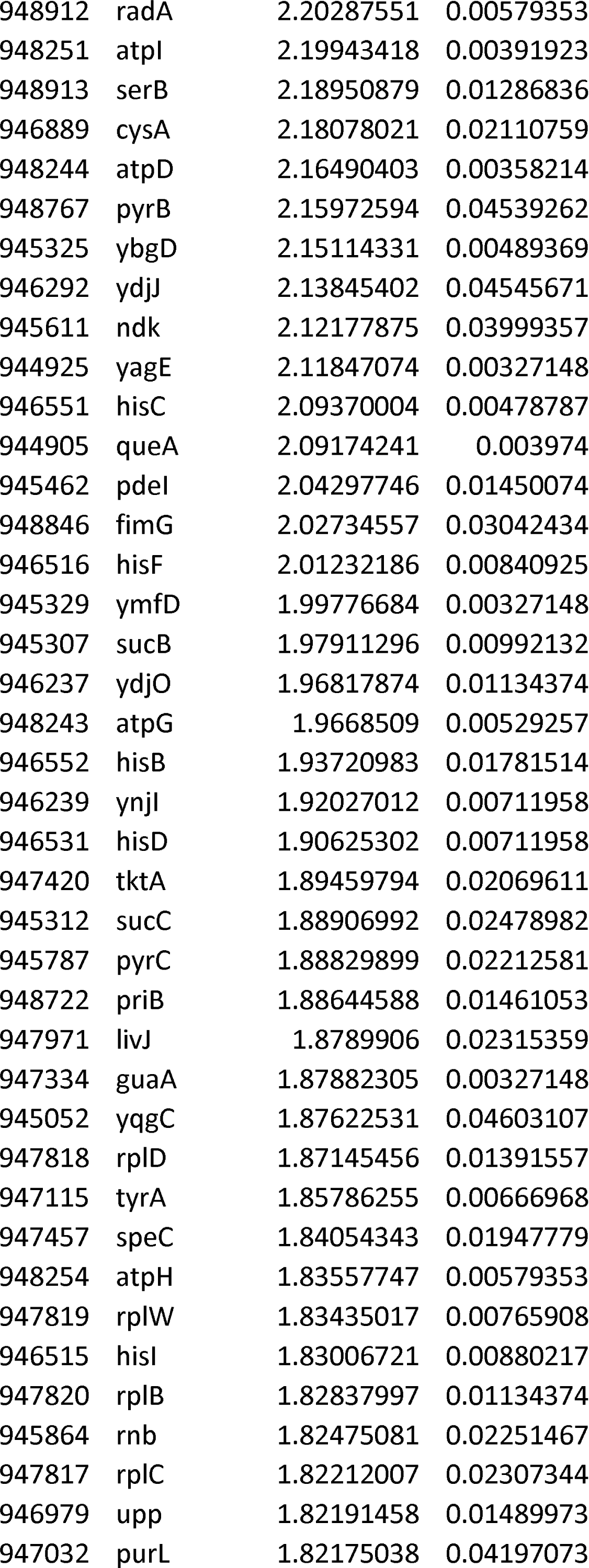
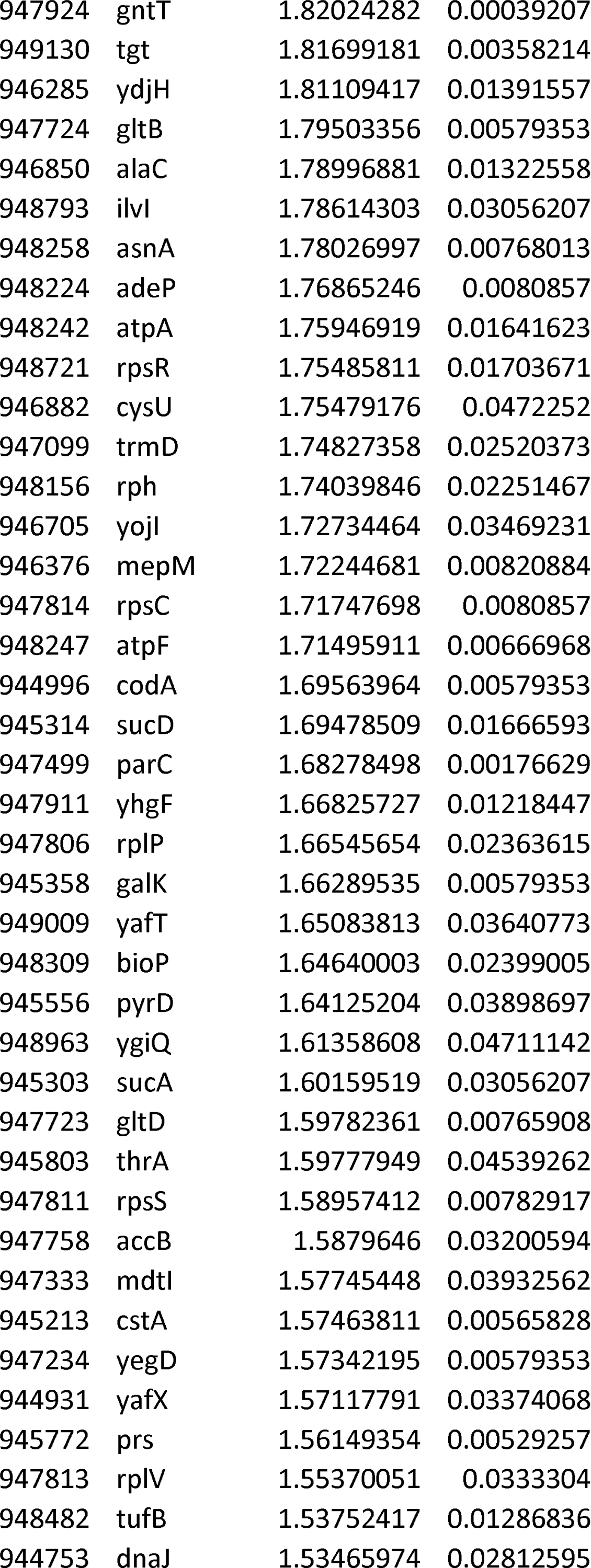
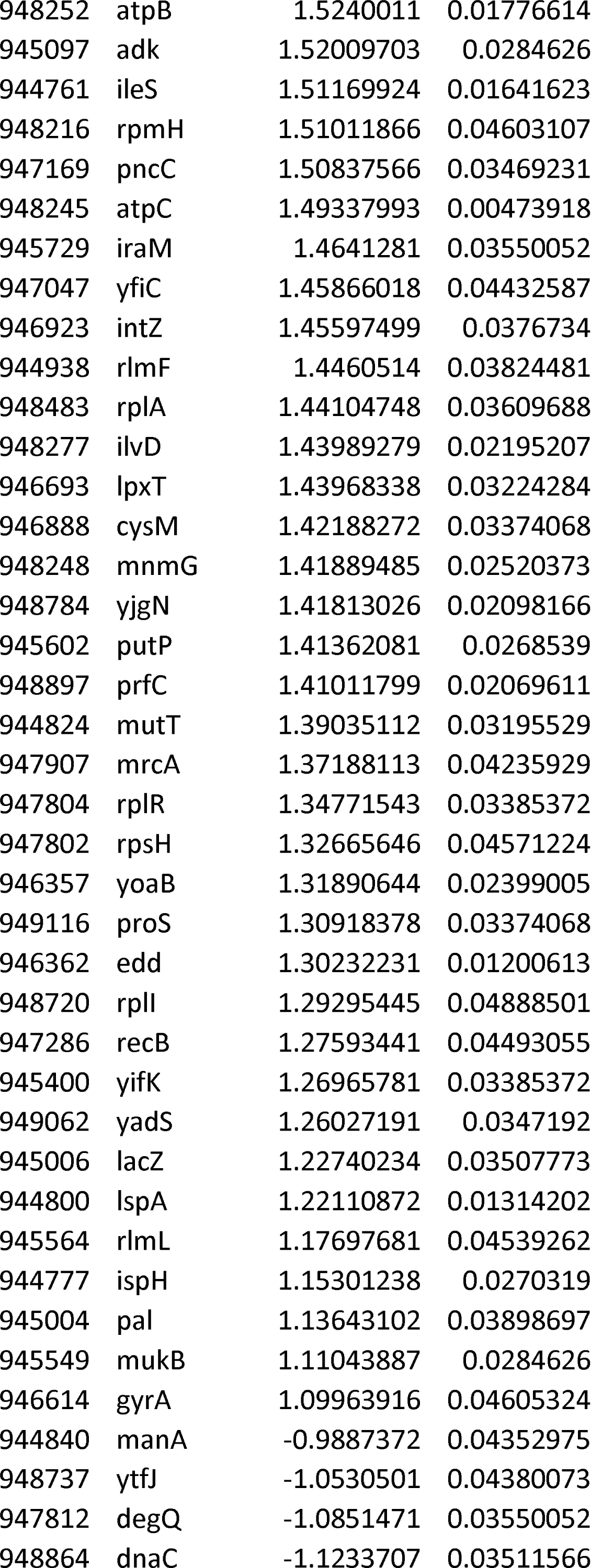
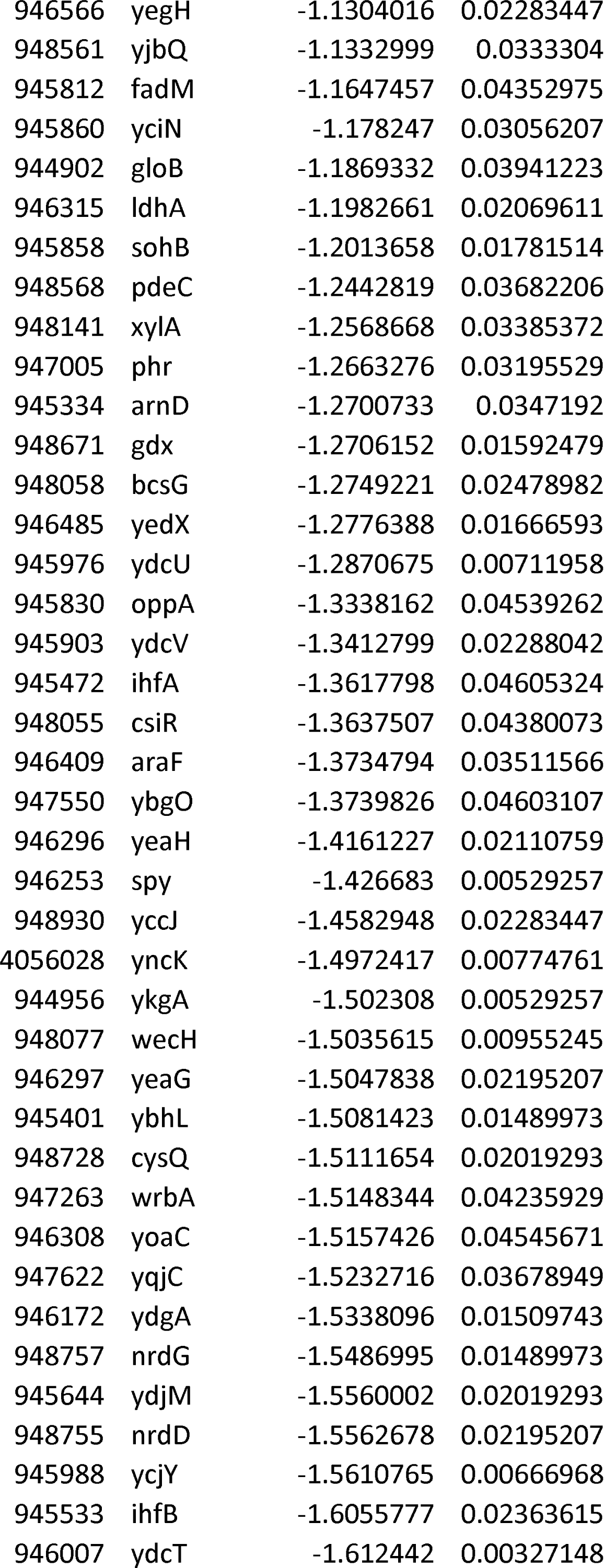
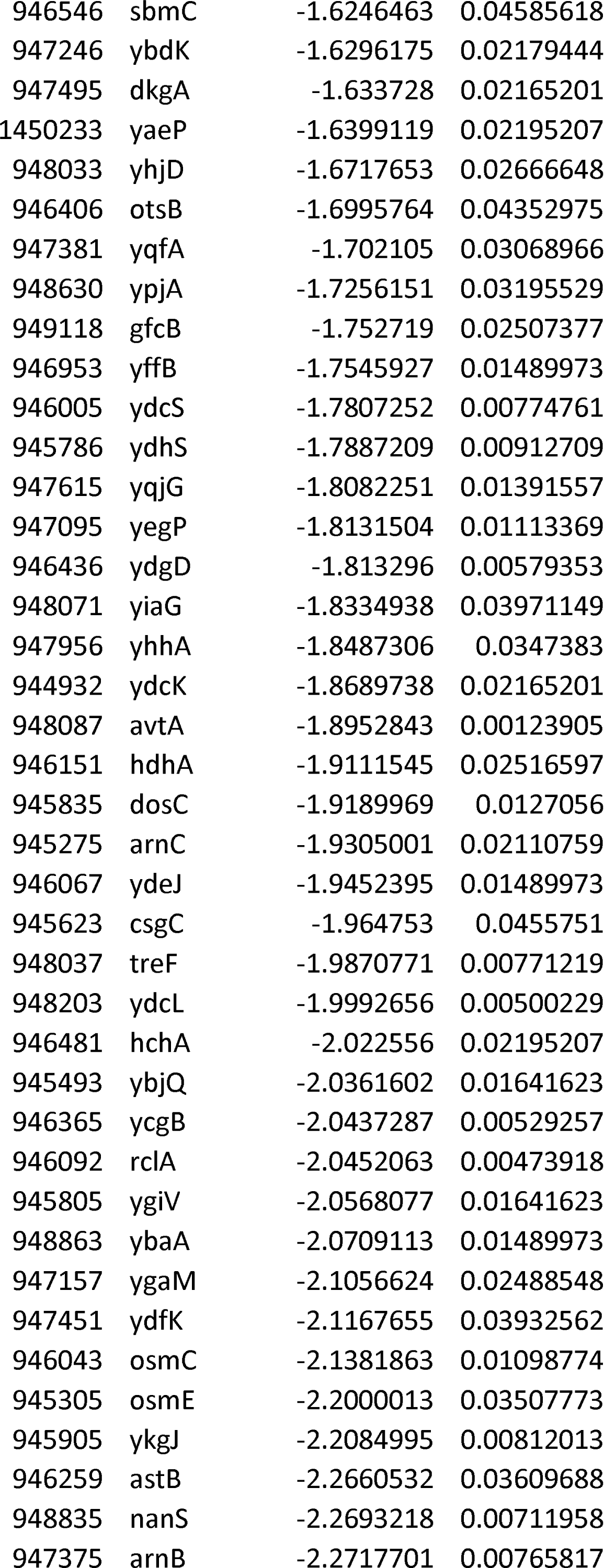
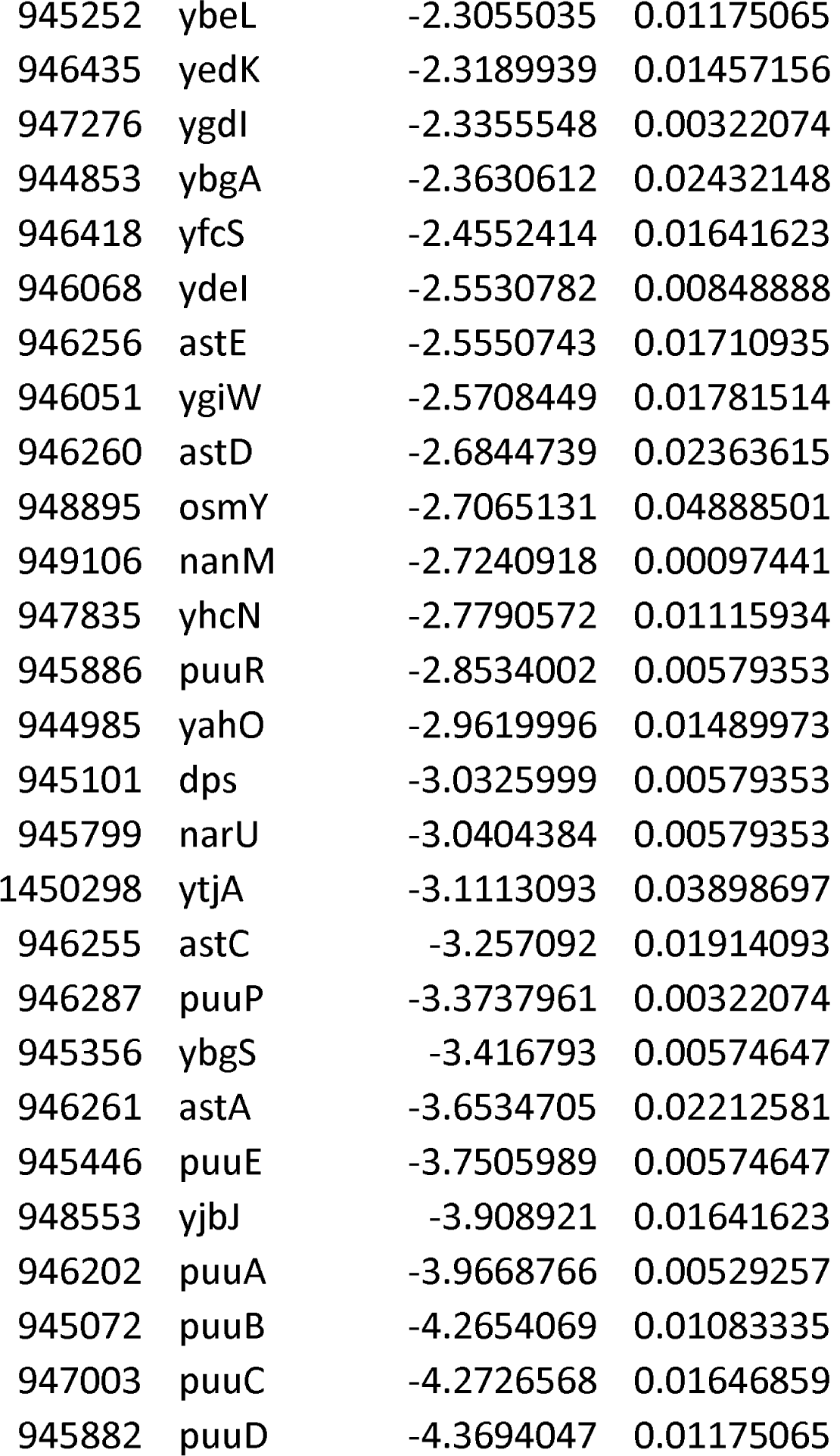
Differentially expressed genes in the strain expressing HUα_2_ P63A relative to the strain expressing wild type HUα_2_

**Table S4.**
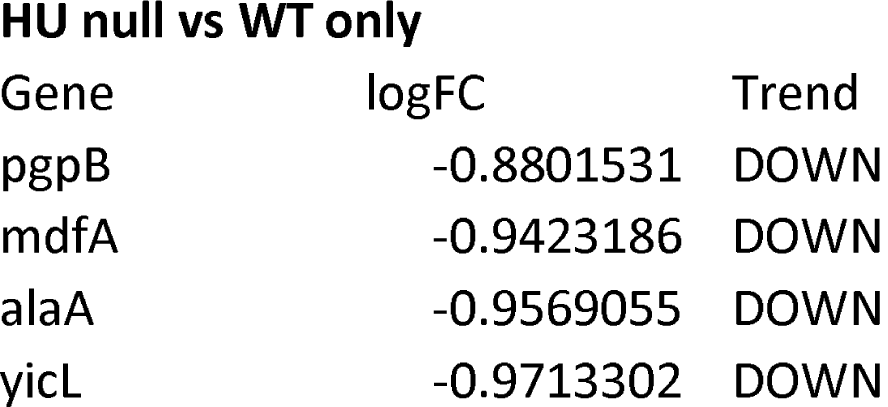
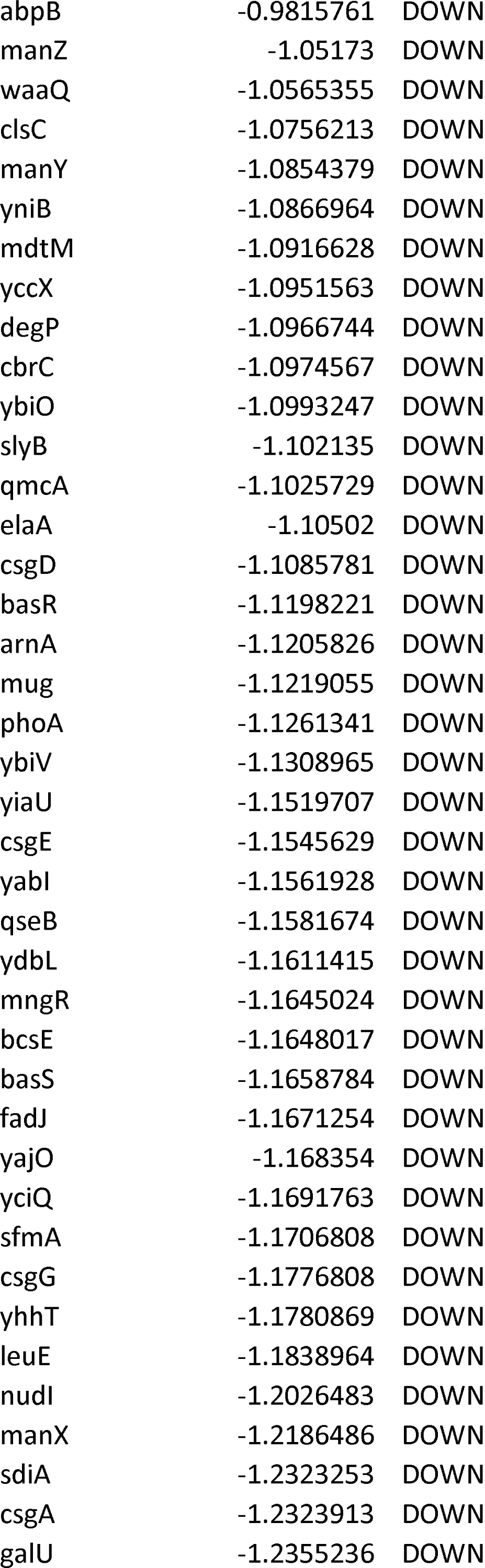
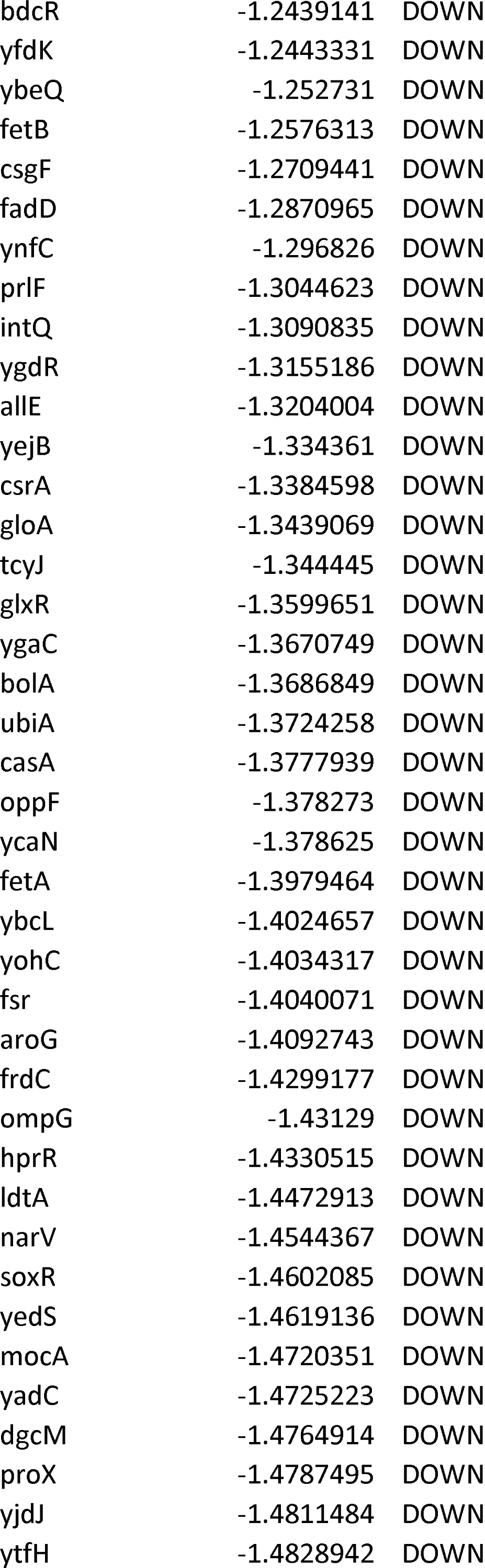
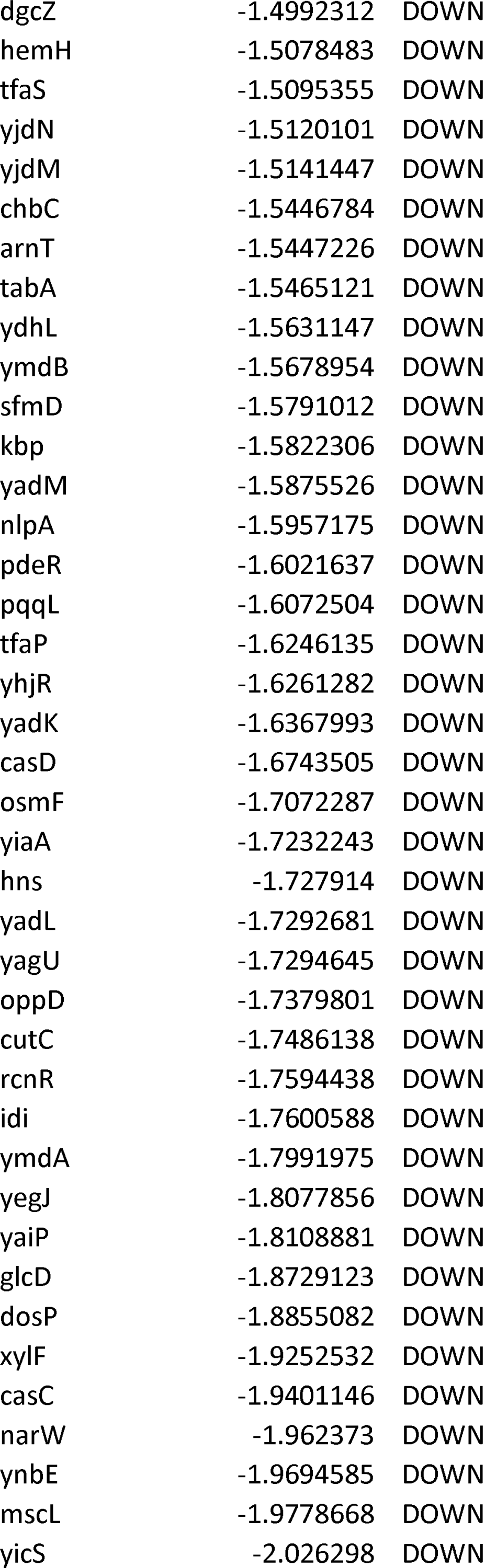
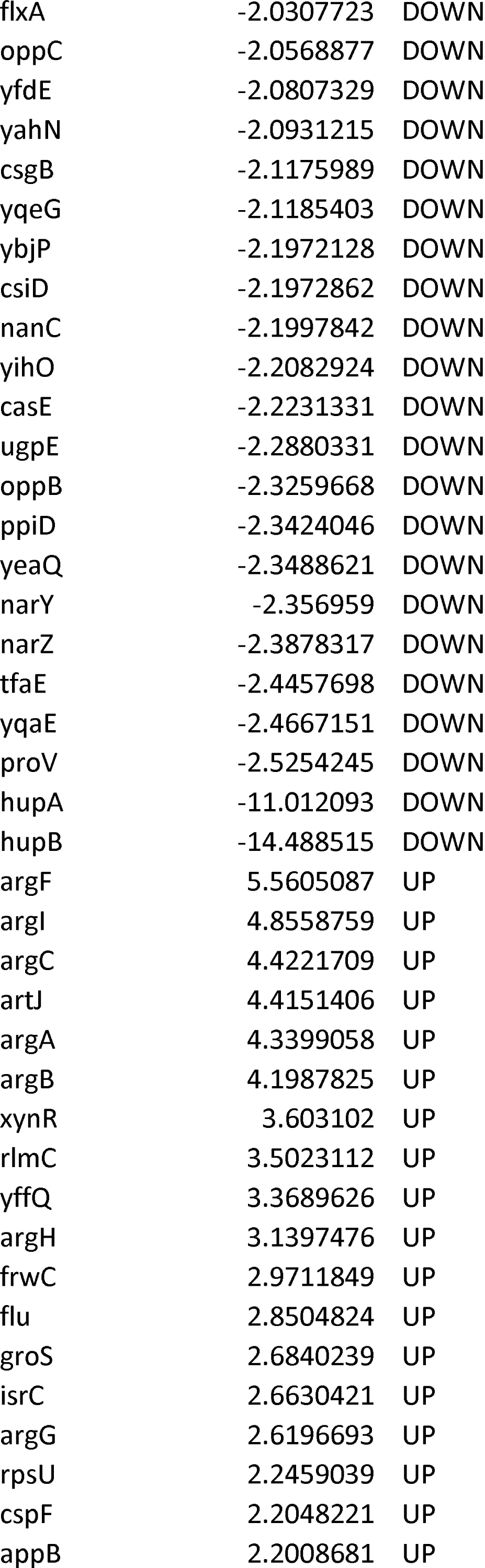
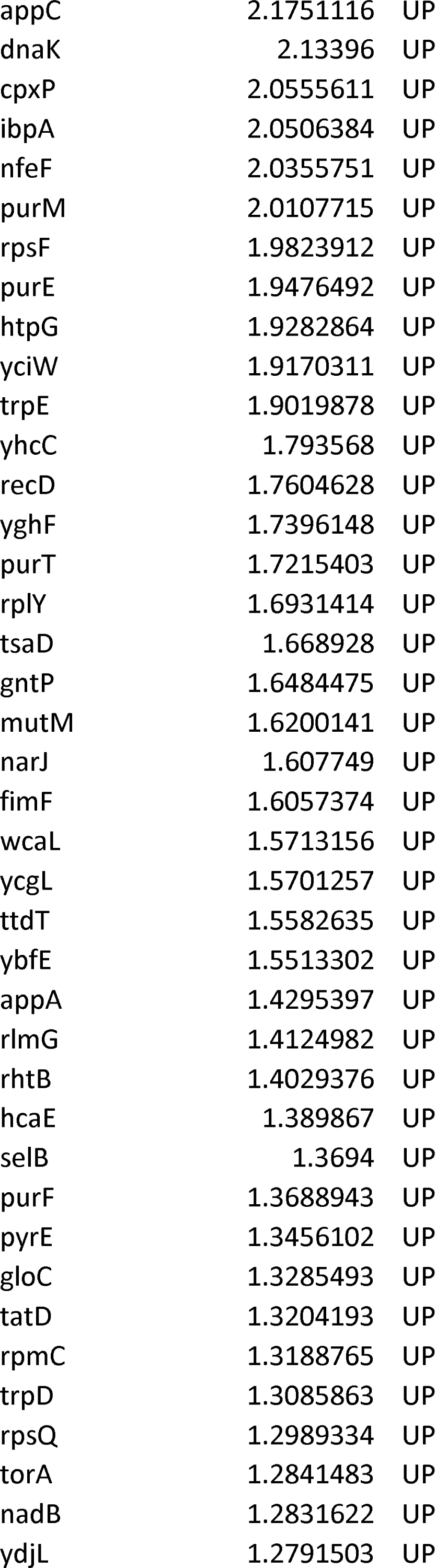
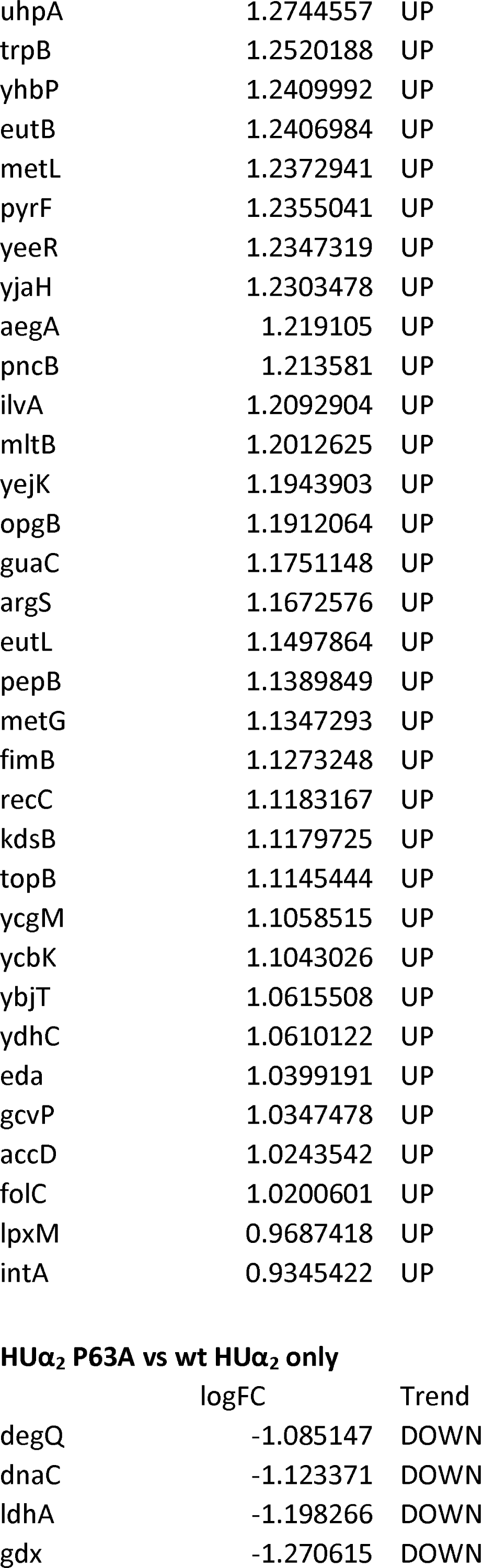
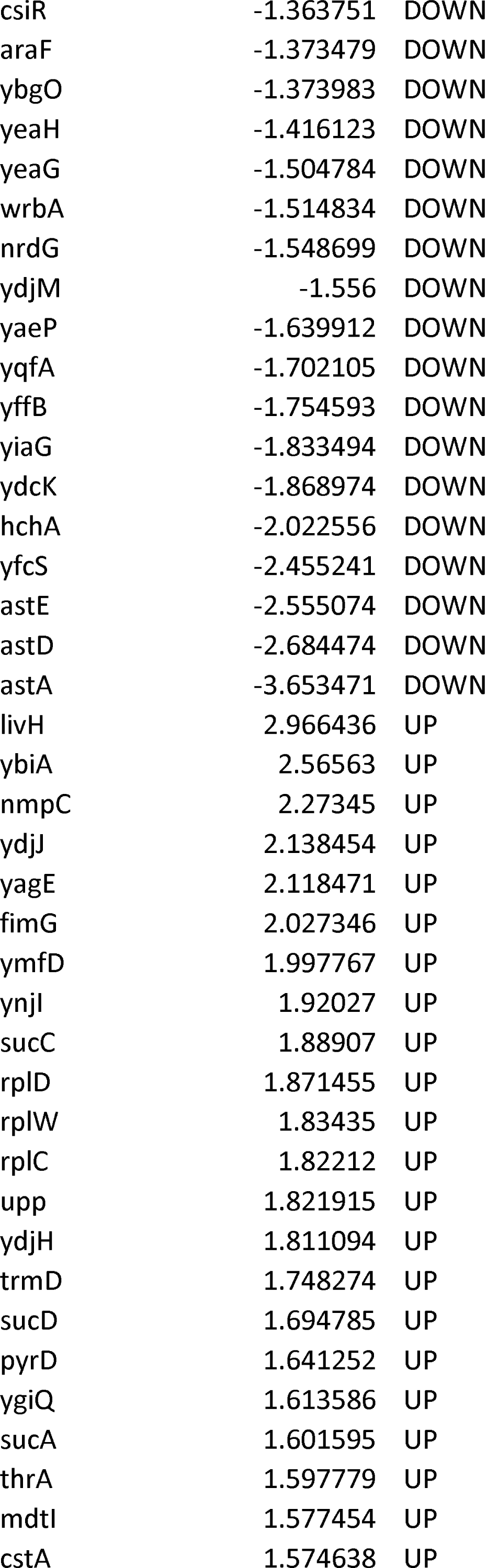
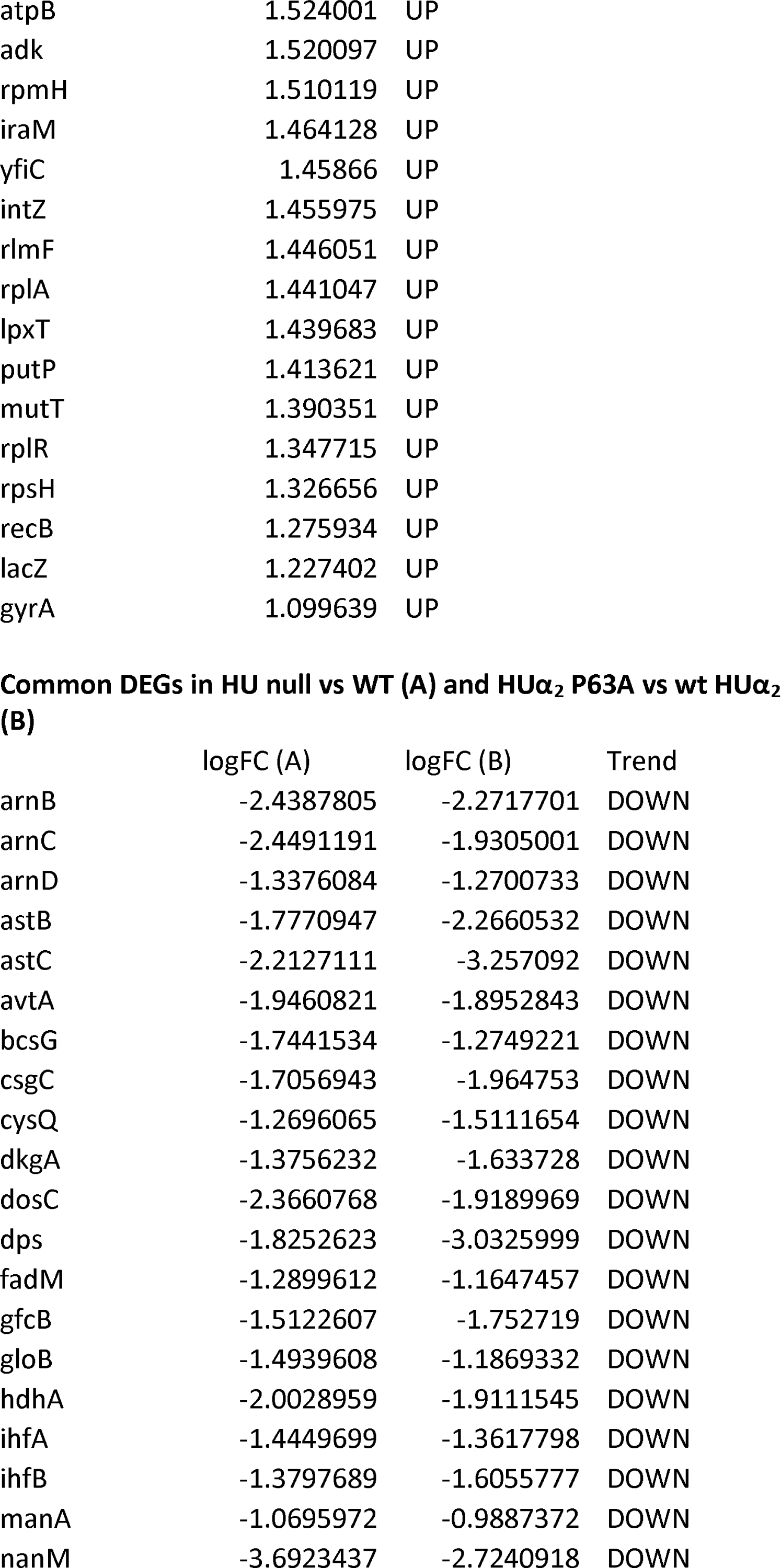
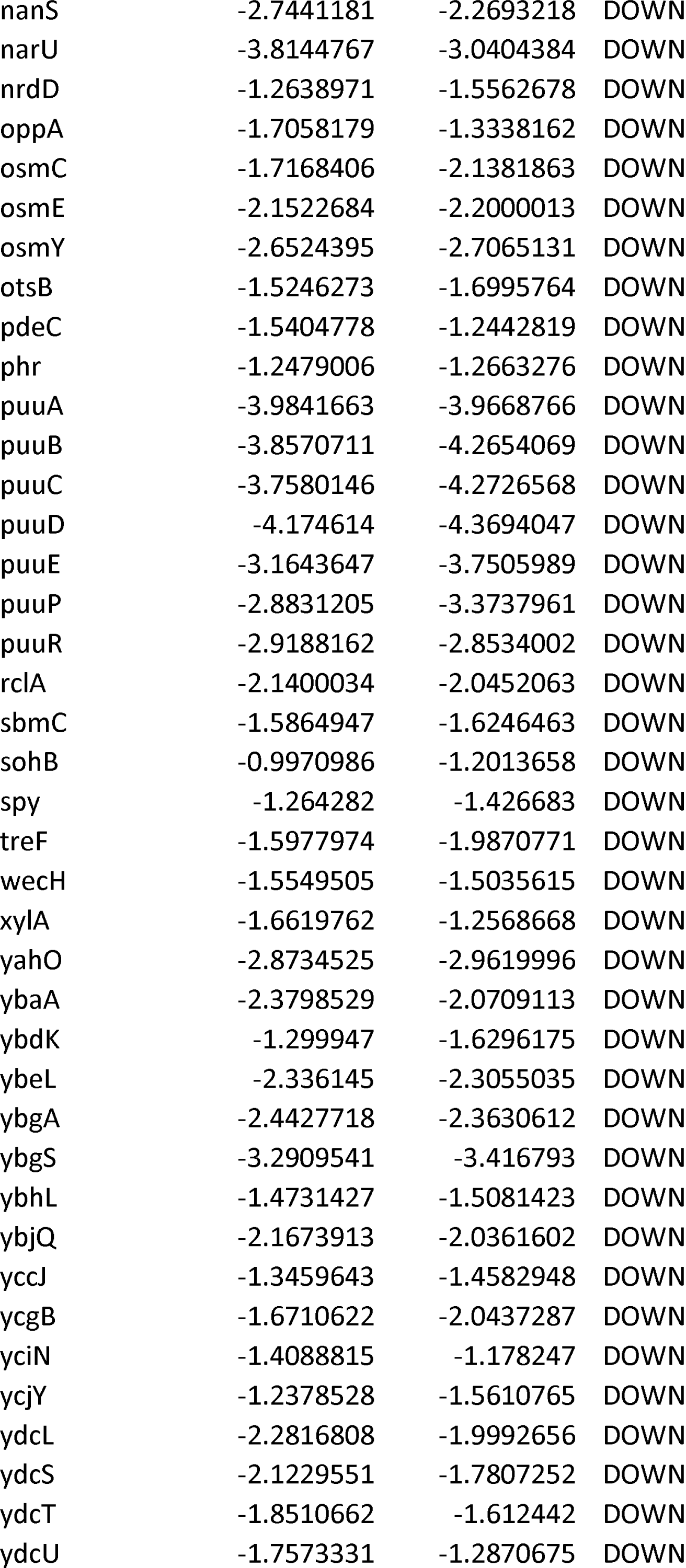
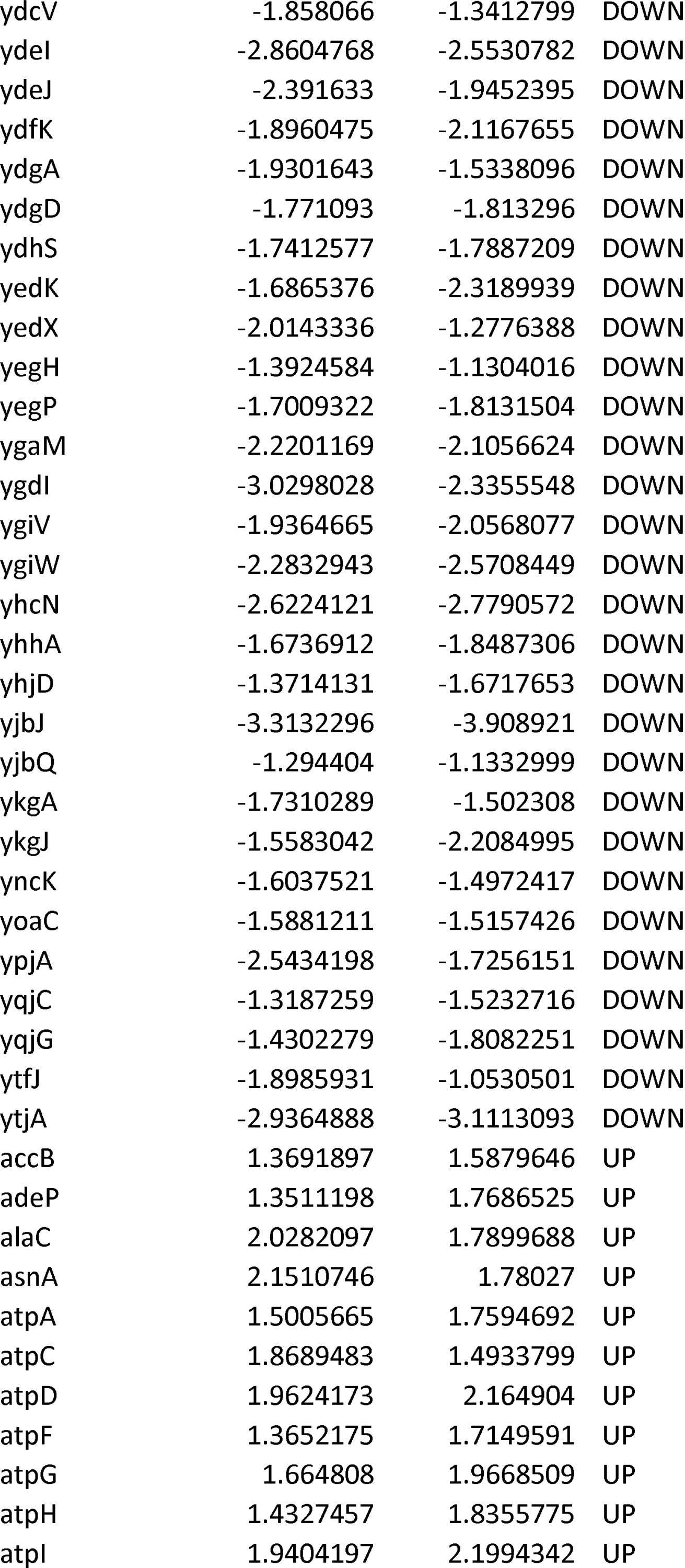
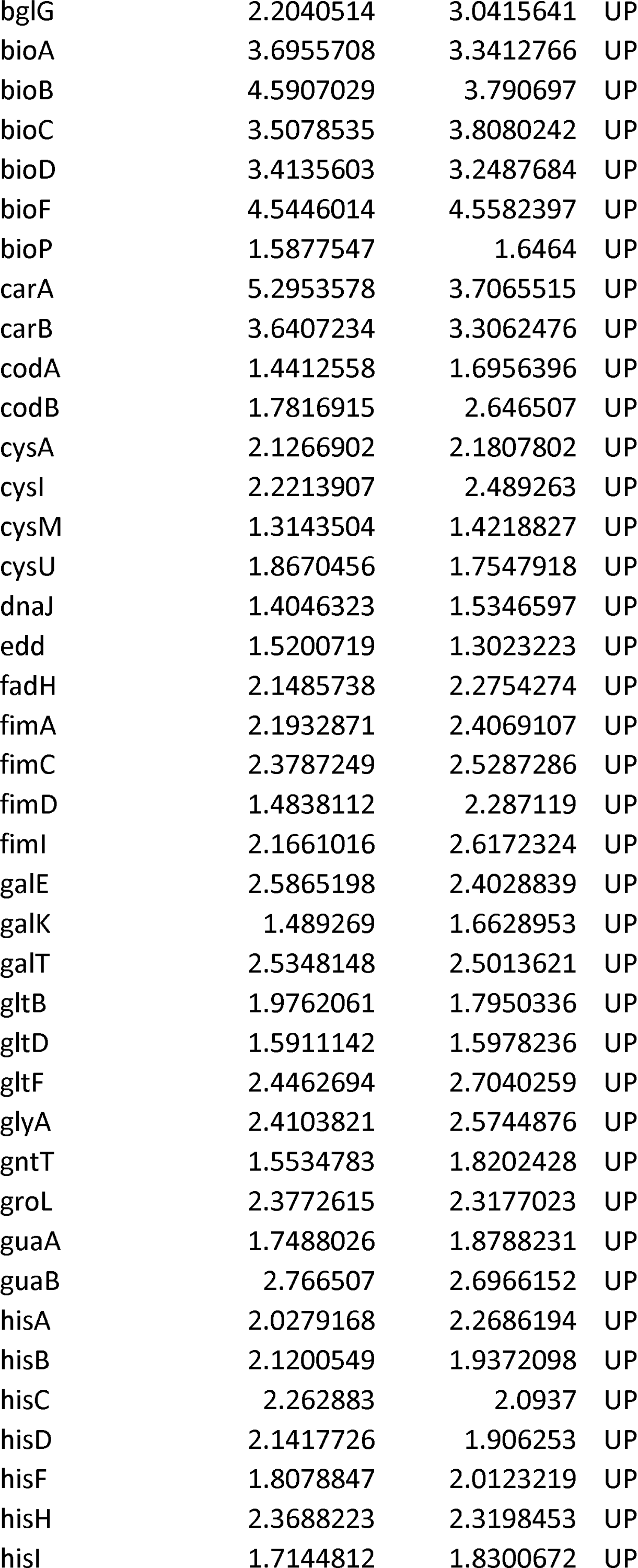
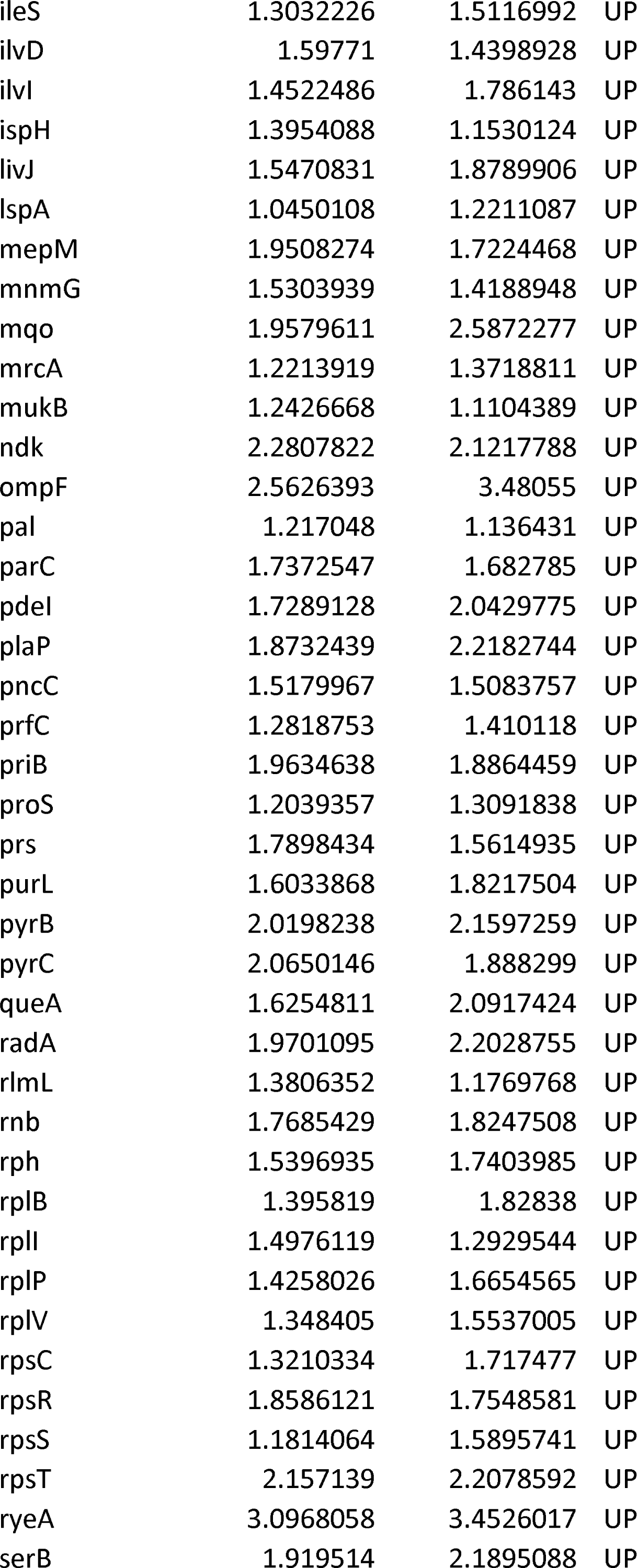
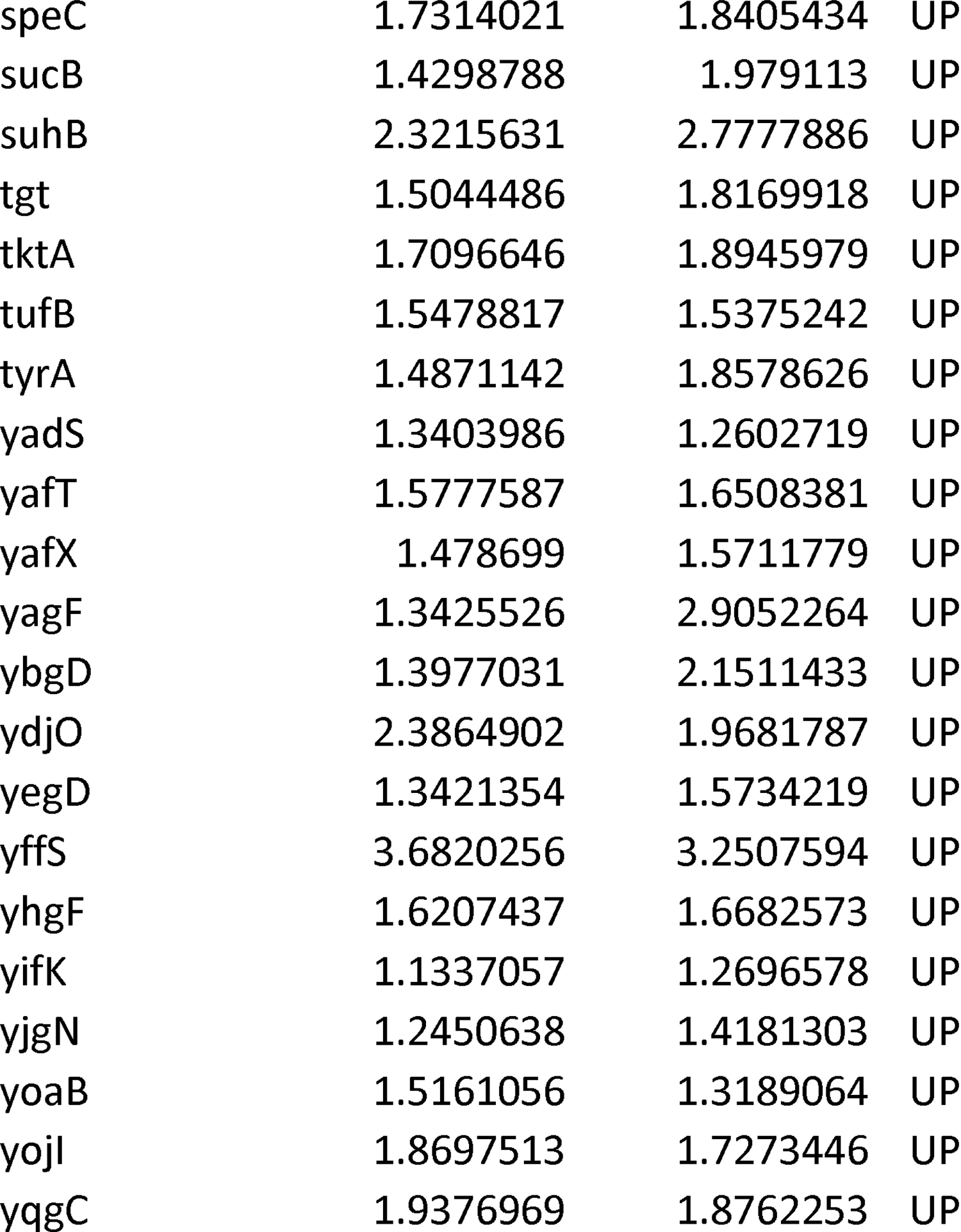
Overlap of differentially expressed genes between the HU null strain (relative to the strain expressing wild type *hup* genes) and the strain expressing HUα_2_P63A (relative to the strain expressing wild type HUα_2_)

